# A fresh perspective on carp feeding behavior in an aquaculture pond and its consequences for individual growth

**DOI:** 10.1101/2025.01.27.635164

**Authors:** Milan Říha, Rubén Rabaneda-Bueno, Marie Prchalová, Lenka Kajgrová, Jaroslav Vrba, Trevis B. Meador, Irina Kuklina, Martin Bláha, Vladislav Draštík, Luboš Kočvara, Lukáš Veselý

**Author notes:** corresponding author **Address of the corresponding author:** Milan Říha Biology Centre CAS, Institute of Hydrobiology Na Sádkách 7 370 05 Česke Budějovice Czech Republic /GSM. The authors had equal contribution (Milan Říha, Rubén Rabaneda-Bueno).

## Abstract

The common carp (Cyprinus carpio) is an important species in global aquaculture. To optimize its production, it is necessary to understand its behaviour in relation to environmental and management factors. This study investigated the spatial and temporal dynamics of carp behaviour in a semi-intensive aquaculture pond under controlled feeding regimes over two growing seasons (2022–2023). Using telemetry and stable isotope analysis, we investigated activity, depth utilisation and feeding behaviour in relation to growth and aquaculture practices.

The activity of carp and their spatial use changed over time, influenced by temperature, food availability and stocking density. Activity peaked in spring and early summer and was highest at night and twilight. In the warmer months, shallow areas (<0.5 m) were favoured, probably due to thermoregulation, oxygen levels or access to prey. Individual variability in foraging site use and food composition correlated with growth, highlighting the importance of behavioural traits in resource acquisition. A flood-induced rise in stocking density in 2022 increased competition for food, resulting in higher activity and lower growth compared to 2023. Stable isotope analysis showed that benthic invertebrates were the main food source, supplemented by cereals, emphasising the importance of a balanced natural and artificial diet.

Our results emphasise the value of telemetry and isotope analysis for improving aquaculture practises. Recommendations include adapting feeding to seasonal and individual behaviour, mitigating density effects and promoting sustainable practises to optimise carp production.

## 1. Introduction

Common carp (*Cyprinus carpio*) is one of the earliest domesticated fish species and remains one of the most popular and commercially significant fish worldwide (Balon, 1995). Due to its economic value, carp has been introduced globally and has become an important species in fish production (Benito et al., 2015; Weber and Brown, 2009). Global aquaculture production of common carp continues to rise, surpassing 4 million tonnes a year, with Europe alone producing 155 thousand tonnes in 2022 (FAO, 2024). The rapid growth in carp production has been driven primarily by the intensification of aquaculture practices over recent decades which, while boosting production, has introduced various ecological challenges (Roy et al., 2020; Verdegem et al., 2023).

Carp farming in Central and Eastern Europe is mainly carried out in artificial ponds that resemble shallow lakes and, over centuries, became an integral part of the region’s landscape and history (Pechar, 2000; Vrba et al., 2024). Moderate-impact or semi-intensive pond management is a common practice of carp farming that depends on both natural food sources (i.e., zooplankton and zoobenthos) and cereal-based supplementary feed (Adámek et al., 2023; Roy et al., 2022). Cereals are estimated to contribute approximately 25–50% to the growth increment of carp (Hlaváč et al., 2014). Yet even when using nutritionally adequate feeds, improper feeding practices can result in significant feed losses that ultimately deteriorate water quality and reduce fish product quality (Hlaváč et al., 2014; Roy et al., 2020). Traditional cereals feeding in carp ponds is often regarded as a significant source of nutrients contributing to pond hypertrophy, with uneaten or poorly digested cereals releasing excess nutrients (Adámek et al., 2023; Roy et al., 2023). Rather than the biological limitations of carp, eutrophication observed in carp ponds has been more attributed to current feeding practices in pond aquaculture (Roy et al., 2020). Therefore, a balanced feeding regime to maximise natural production and optimise artificial feeding is key to maintaining high carp production while increasing economic efficiency and reducing the ecological footprint of carp farming (Hlaváč et al., 2014; Roy et al., 2022). Addressing these challenges and improving feeding efficiency requires a thorough understanding of carp foraging behaviour, which depends on intrinsic factors (e.g. biological and physiological needs) and extrinsic factors (e.g. environmental conditions and food availability) (Kajgrová et al., 2024; Orság et al., 2023; Potužák et al., 2007; Roy et al., 2022; Stanivuk et al., 2024).

Previous studies have demonstrated that carp exhibit significant dietary plasticity, particularly favoring benthic macrozoobenthos over zooplankton as they mature (García-Berthou, 2001; Rahman and Meyer, 2009). In semi-intensive aquaculture systems, however, cereal-based supplementary feed plays an important role in both carp nutrition and overall pond ecosystem functioning (Roy et al., 2022). Feeding cereals satiates carp and spares natural food from being overgrazed, additionally high protein sparing potential and low digestible phosphorus-to-protein ratio in cereals help to spare the protein and phosphorus from natural prey. While cereals supply energy to carps, the natural food (i. e., zooplankton and zoobenthos) remains essential as a source of key amino acids, fatty acids and to support the digestion of phosphorus from the provided supplementary feed (Roy et al., 2024). A lack of such essential food sources, which cannot always be substituted, can impair individual fitness and growth (Martin-Creuzburg et al., 2008; Schälicke et al., 2019). This highlights the necessity of balancing artificial and natural food sources to optimize carp growth, sustain ecosystem health, and ensure the long-term sustainability of aquaculture practices.

One approach to reveal the mentioned balance is research on carp movement within the pond system. Advances in tracking technologies offer valuable insights into carp behavior in aquaculture systems (Nathan et al., 2022; Özgül et al., 2024). Modern methods such as telemetry, enable precise monitoring of fish movements and interactions, aiding in the optimization of feeding strategies, waste reduction, and improved productivity (Özgül et al., 2024). Previous telemetry studies conducted in ponds with supplementary feeding indicate that carp intensively utilized feeding zones while also exploring other areas of the waterbody during feeding periods (Jurajda et al., 2016). This behavior allows them to access both supplementary food and the natural prey. However, there is limited information on how fluctuations in carp’s space utilization are influenced by factors such as food availability, temporal changes, temperature, initial fish density, presence of invasive species or competition, and how these developed over the year and/or production cycle.

The choice of feeding regime plays a critical role in aquaculture (Attia et al., 2012) and diel feeding rhythms are particularly important for carp foraging (Klaren et al., 2013). While laboratory experiments have shown that carp preferred nocturnal feeding when given the opportunity to feed on demand (Klaren et al., 2013), field studies presented more ambiguous results. Some studies report diurnal foraging behavior (Rahman and Meyer, 2009), while others observe mainly nocturnal activity (Bajer et al., 2010; Benito et al., 2015; Hundt et al., 2022; Žák, 2021). Additionally, individual carp may exhibit distinct daily activity patterns, indicating a level of behavioral plasticity (Benito et al., 2015; Žák, 2021). Understanding these diel variations is essential for improving aquaculture practices. Currently the procedures are based on workers’ schedules with only limited respect to fish foraging habits. Aligning feeding schedules with the natural or preferred foraging times of carp would enhance production via increased feeding efficiency and faster growth rates while reducing waste.

Important features of the foraging behaviour of carp are also linked to their social, cognitive and individual traits. They demonstrate rapid learning, recognizing feeding sites and adapting to feeding schedules (Bajer et al., 2010; Ghosal et al., 2018; Zion et al., 2007). Individual differences, including personality traits, further influence feeding success, with bolder carp more likely to explore new sites (Górecki et al., 2019; Huntingford et al., 2010). Larger, socially dominant individuals often monopolize feeding areas, which correlates to higher muscle fat content and body condition (Hundt et al., 2022; Jurajda et al., 2016). These findings underscore the importance of individual traits and social dynamics in shaping carp foraging behavior, with implications for aquaculture practices.

A key approach to understanding the utilization of artificial and natural food sources in carp production is stable isotope analysis (SIA). This method enables the identification of food source origins and is widely applied in fish biology and aquaculture studies. For example, Binhe Gu et al. (1996) examined trophic overlap between two planktonivorous fish in ponds, while Ray et al. (2017) investigated the contributions of clear-water recirculating aquaculture systems (RAS) and biofloc-based systems to shrimp production (*Litopenaeus vannamei*).

Similarly, SIA has been used to trace the origin of fish production—whether from aquaculture or natural habitats (Gamboa-Delgado et al., 2014). When combined with research on the spatial feeding habits of carp in ponds, these methods may provide valuable insights that support the development of efficient, sustainable, and welfare-focused aquaculture practices.

In this study, we investigated the behavior of carp in a semi-intensive aquaculture pond, focusing on their spatial and temporal dynamics under a controlled feeding regime. Our objectives included examining seasonal and diel patterns in carp space use and activity within the pond, as well as exploring individual variation in feeding behavior and its implications for diet composition and growth. We hypothesised that: i) carp show significant temporal shifts in activity and horizontal and vertical space utilisation; ii) these shifts reflect the feeding regime and changes in environmental conditions in the pond; iii) significant individual differences in activity and feeding area utilisation would lead to measurable differences in growth rates and diet composition. To test these hypotheses, we tracked carp behavior over two growing seasons using a fine-scale telemetry system. Additionally, stable isotope analysis was employed to evaluate diet composition. Further, seasonal assessments of natural food availability were conducted to record natural prey supply and to understand its influence on carp behavior.

## 2. Material and methods

### 2.1 Study site and pond management

The study was conducted in the small aquaculture pond Louzek, located in Central Bohemia, Czech Republic (GPS coordinates 49°32’34.43"N, 13°54’56.10"E; Fig. 1), over two growing seasons in 2022 and 2023. Pond Louzek has an area of 1 hectare, a maximum depth of 3 meters near the dam, and an average depth of 1.5 meters. Water transparency (Secchi depth) ranged from 1.4 meters in May to 0.4 meters in October. Mean total phosphorus (TP) concentrations were 0.19 µg/L in 2022 and 0.17 µg/L in 2023. The pond lacks underwater structures, with the reed (*Phragmites australis*) covering approximately 13% of the shoreline to a depth of 1 meter along the western part of the pond. Additionally, Glyceria (*Glyceria sp.*) covers about 3% of the shoreline in the northeastern corner (Fig. 1). Before the experiment, the pond was drained in autumn and left unfilled for several months. The harvesting area and filling channels were limed prior to refilling. It is supplied by a small stream within the Skalice River catchment.

**Figure 1.**
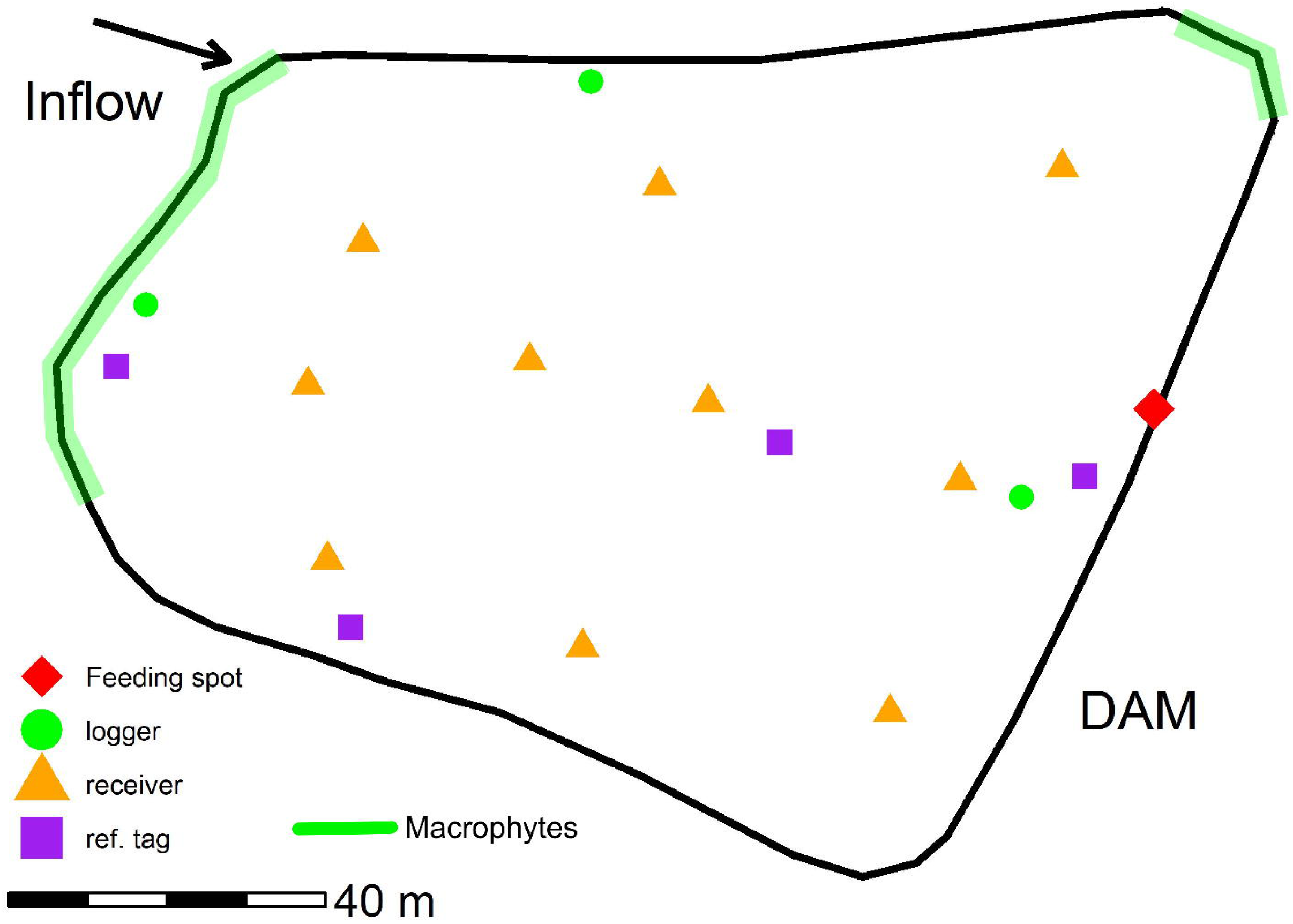
Map of the study area with the most important features (feeding site and water macrophytes distribution) and locations of the telemetry system.

In 2022, a total of 772 carp individuals (mean body weight: 202 ± 56 g; total biomass: 156 kg) were stocked in the Louzek pond. In 2023, 830 carp individuals (mean body weight: 171 ± 46 g; total biomass: 142 kg) were stocked.

In June 2022, heavy rainfall caused local flooding, washing numerous small fish from upstream ponds into the Louzek pond. This event introduced additional fish into the pond environment, which influenced the observed biomass and species composition in 2022 (see chapter 3.6).

Feeding was carried out at a point near the center of the dam (Fig. 1). The fish were fed with cereals (triticale) following established practices (Füllner, 2015). Feeding took place from 26 June to 8 September 2022 and from 4 July to 12 September 2023. The fish were fed at intervals of 2 to 7 days (mean 4.7 ± 1.6 days), reaching 17 and 16 feedings in 2022 and 2023, respectively. The dose for one feeding was 50 or 100 kg, reaching a total of 1050 and 1150 kg of cereals in 2022 and 2023, respectively.

### 2.2 Tagging and telemetry tracking

#### 2.2.1 Tagging

A total of 36 carp individuals (20 in 2022 and 16 in 2023) were tagged for this study. All fish were two years old (K2) and originated from the “Blatenská Ryba Ltd.” The mean total body length and mass were 308 mm/438 g in 2022 and 319 mm/552 g in 2023, respectively (see Table 1 for additional details).

**Table 1.** Summary of biometric data for tagged individuals in 2022 and 2023. The table contains the number of tagged individuals (N), the mean total length (mm), the minimum and maximum total length (mm), the mean weight (g), the minimum and maximum weight (g) and the number of recovered individuals during harvest each year.

| Year | N | mean<br>TL<br>(mm) | min TL<br>(mm) | max<br>TL<br>(mm) | W (g) | min W<br>(g) | max W<br>(g) | N<br>recovered |
| --- | --- | --- | --- | --- | --- | --- | --- | --- |
| 2022 | 20 | 307.6 | 265 | 362 | 438.4 | 310 | 722 | 18 |
| 2023 | 16 | 318.8 | 280 | 385 | 552.1 | 338 | 1072 | 11 |

Prior to tagging, each carp was anesthetized using 2-phenoxy-ethanol (SIGMA Chemical Co., USA) at a concentration of 1 ml Lu¹ (Kolářová et al. 2007). Fish were then measured, weighed, and prepared for tagging. A 1.5–5 cm incision was made along the ventral surface, just posterior to the pelvic girdle, to insert the transmitter (Lotek Wireless Inc., MM-M-8-SO-TP, 42 × 8.5 mm, 6 g in air, equipped with depth and temperature sensors, burst rate: 60 s). Tag weight ranged from 0.5% to 1.9% of the fish’s body weight (Table 1). Antibiotics (Betamox LA, Norbrook Manufacturing Ltd, Ireland) were applied to the body cavity to prevent infection. The incision was closed using several independent sutures (Polyamide monofilament, Resolon 3EP 2/0USP, Resorba, Czech Republic).

After tagging, carp were bathed briefly (10 seconds) in a potassium permanganate solution to prevent infection and then placed in fresh water for full recovery. The tagged fish were subsequently held at the aquaculture facility for post-tagging monitoring: 20 days in 2022 and 16 days in 2023. During this period, the carp were kept in a pool (pool dimensions 2.70 x 1.6 x 1 m with water level 0.65 m and constant water run) for observation of potential post-tagging mortality. Two individuals died in 2022 and were replaced with new fish, while no mortality was observed in 2023. The tagging was carried out on 19 April 2022 and 18 April 2023, the fish were released on 9 May 2022 and 4 May 2023. The pond was completely fished out and all fish that survived were recaptured on 1 November 2022 and 20 October 2023. During harvest, surviving tagged fish were euthanized, measured (to the nearest mm), weighed (to the nearest gram), and muscle samples were collected for stable isotope analysis.

#### 2.2.2 Telemetry tracking

A MAP positioning system (Lotek Wireless Inc., Canada) was deployed in the pond. The systems consisted of 10 receivers (Lotek Wireless Inc., WHS3250) deployed in arrays with distances between pairs of receivers ranging from 14.5 to 101.2 m (mean 54.2 ± 23.5 m) (Fig. 1). The exact position of deployed receivers was measured using a high-precision GNSS-unit (Spectra Precision, Promark 220, USA). Depth of receivers varied between 1 -1.5 m. According to range testing done prior to the study, in this pond (July 2020), such settings of receiver arrays provide full coverage in the whole pond. The systems were installed in both years from the beginning of April to mid-October and the data was downloaded manually every two months.

The accuracy of the system was verified using 5-6 stationary reference tags (MM-M-16-50-TP, burst rate 25 sec, Lotek Wireless Inc., Canada) placed at 4 locations in each lake (near the dam in the center of the pond in open water at a depth of 0.5 and 1.3 m and at 2 nearshore locations at a depth of 0.5 m). In addition, the accuracy was tested using reference tags that were pulled under a boat in June 2022 and October 2023 (see details in the Supplementary material 1).

### 2.3 Food availability and water temperature sampling

Zooplankton and zoobenthos, as a key natural food sources, were sampled in both seasons to assess temporal changes in their availability.

This study focused on the total density of larger zooplankton (>200 µm), as adult carp can efficiently retain large food sources (>250 μm) with their branchial sieve (Sibbing et al., 1986). Mixed water samples from the surface layer were collected using a 1 m-long van Dorn sampler (6.4 L volume) and integrated into 50 L washed plastic containers by sampling at seven points along a linear transect in the open water zone of the pond. A subsample was filtered through a 200-µm mesh and stored in a plastic bottle for further laboratory analysis. Additional details on zooplankton processing are provided in (Vrba et al., 2024).

Macrozoobenthos for biomass analysis was collected from the pond bottom at 10 randomly distributed sites using an Ekman-Birge sampler. The samples were preserved in 96% ethanol and later identified to the lowest possible taxonomic level in the laboratory.

In order to assess the carp’s preference for certain food sources and to take into account potential variations in stable isotope signals within the pond, zooplankton and zoobenthos, terrestrial food sources (hereafter referred to as TF sources -including emergent aquatic insect imagos, terrestrial insects, and spiders and cereals were included as a putative food sources. For stable isotope analysis, zooplankton was collected using a zooplankton net with a mesh size >250 µm from at least three distinct locations around the pond’s bank, with a minimum of five tows at each location. Zoobenthos, were collected using the Ekman-Birge sampler. Organisms from five samples were combined into three composite subsamples. TF sources were collected simultaneously using sweep netting along the pond bank (0–2 m from the shoreline). Sampling of these food sources for stable isotope analysis were performed at the end of August and September.

To obtain abiotic parameters that may influence the spatial distribution and activity of the carp, we monitored the water temperature throughout the pond. Temperature data were collected using eight data loggers (Onset, USA, HOBO Pendant temp/light 64K) positioned in three locations to cover both the open water and the near shore zones (Fig. 1). Near the dam, data loggers were attached at 0.5 metre intervals along a rope that reached from the surface to a depth of 2.5 metres. This rope was connected to a floating buoy anchored to the bottom of the pond so that the vertical temperature profile in the open water could be monitored continuously. In the nearshore area, the data loggers were positioned at a depth of 0.5 metres at the bottom of the pond and attached to a brick to ensure stability. This arrangement, with a 5-minute measurement interval, provided a high spatial and temporal resolution for monitoring temperature variations across both open water and shallow areas.

### 2.4 Stable isotopes analysis

Before analysing the stable isotopes, all samples (fish and food sources) were lyophilized for 48 hours and ground to a fine, homogeneous powder. Approximately 0.6 mg of organism-based tissue and 1 mg of plant-based tissue were weighed into tin capsules using an electronic microbalance. The analyses were performed using a Carlo Erba Flash EA 1112 elemental analyser connected to a Thermo Finnigan DELTAplus Advantage continuous-flow isotope ratio mass spectrometer (Thermo Electron Corporation, Waltham, MA, USA).

Vienna Pee Dee belemnite and atmospheric N2 were used as reference standards for δ13C and δ15N, respectively. To control instrument stability, muscle of the Northern pike (Esox lucius) of known isotopic composition was analyzed every six organism tissue based samples. Similar approach were applied for plant based samples using mate powder as internal standard. Results are expressed using the conventional δ notation as per mil (‰) difference from the international standards. The analytical precision was < 0.1 ‰ for δ13C and < 0.3 ‰ for δ15N.

### 2.5 Data analysis

#### 2.5.1 Telemetry data

Individual fish locations were initially calculated using the manufacturer’s proprietary positioning software, UMAP v.1.4.3 (Lotek Wireless Inc., Canada). Fish depth was recorded by the tag’s internal sensor, which has a resolution of 0.2 m. We visually location estimates, along with the depth profiles for each fish. Instances where both horizontal and vertical positions remained constant, indicating no movement, were interpreted as either a deceased fish or an expelled tag. This was the case for five fish in 2023.

In 2022, extensive tag malfunctions were observed, with only six tags remaining functional until at least mid-August, while the rest failed by June for unknown reasons. In 2023, only one tag experienced a similar malfunction. Consequently, no movement data was available for the fish with malfunctioning tags. In summary, due to tag malfunctions and fish mortality, we obtained telemetry data for six individuals in 2022 and ten individuals in 2023. However, we recaptured 18 out of 20 tagged individuals in 2022 and 11 out of 16 in 2023. For these recaptured fish, growth and stable isotope data are available.

To estimate the space use of individual fish, we employed the autocorrelated kernel density estimation (aKDE) method using the ctmm R package (Calabrese et al., 2016). This method accounts for temporal autocorrelation in movement data, providing unbiased utilization distributions (UDs). For each individual, a variogram analysis was conducted to quantify movement autocorrelation, and an appropriate movement model was selected based on Akaike’s Information Criterion (AIC). The selected model was used to compute individual UDs with aKDE. To estimate population-level space use, we applied the population kernel density estimation (pKDE) approach, which combines individual UDs into a single population-level UD. This method integrates information from all individuals while accounting for their spatial distribution and temporal sampling effort. The aKDE and pKDE were computed for each month to analyze temporal variations in individual or population-level space use, with 95%, 50%, and 25% isopleths extracted to represent individual or the overall population’s home range, core area, and intensive use zones.

Activity of fish was calculated as swimming speed (expressed in meters per second, m.sec^-1^) between two consecutive locations using true 3 dimensional distance (Lennox et al., 2024). Depth of fish was measured by an internal tag sensor and transmitted together with a tag identification.

Utilization of the feeding ground: We considered an individual to be at a feeding site if they were within 15 m of a feeding site, and we calculated the daily proportion of locations at the feeding site (FGU) by dividing the number of individual locations at the feeding site by the total daily number of individual locations.

### 2.6 Statistical analysis

### 2.6.1 Fish activity

Swimming speed (“ss_3d”) was analyzed as a function of date, hour, body weight and average temperature across years. To account for non-linear relationships and temporal dynamics, generalized additive models (GAMs) were implemented using Bayesian analysis with the R package brms (Bürkner, 2017), which provides a flexible framework for handling hierarchical data while incorporating smooth terms and dealing with uncertainty -conditions that were met by the imbalance of data between years in our study. Null models with different distribution families, including Student’s t, Gamma and Inverse Gaussian, were compared to determine the best fit for the swimming speed data. Separate models were fitted for each year, allowing independent estimation of year-specific temporal and environmental effects while accounting for individual variability. The models from each year were compared using leave-one-out cross-validation (LOO) (function loo_compare() in brms), and the model with the highest expected log predictive density (elpd) and smallest standard error difference was selected for further analysis. The gamma distribution family with a log link was the best fit in both cases, as it provided the best balance between flexibility and fit to the positively skewed values of swimming speed. Once the distribution family was selected, more complex models with different predictors and interaction terms (e.g. tensor product smoothing) were fitted. Each model included smooth terms for date, for the effects of fish weight and temperature, and an interaction between date and time of day (hour) using the smooth tensor product (t2(date, hour), with the spline basis used in this case containing three basis functions to reduce complexity and capture important trends. To account for the cyclical nature of time, a cyclic spline was applied to the hour variable, while the date was adjusted with a thin plate spline (bs=c("tp", "cc")). Random intercepts for individual fish (fishID) were included to account for repeated measurements and individual variability. We used the default priors provided by the brms package as these were deemed appropriate given the lack of prior knowledge of the parameters. The models were fitted with four chains of 2000 iterations each (including 1000 iterations for the warm-up phase) to ensure robust posterior estimates. Posterior predictive checks and diagnostic metrics such as effective sample size and R-hat values were used to assess model fit and convergence. Conditional effects plots were used to visualize the relationships between swimming speed and predictors. The final selected models are represented by the same formulas, as they are structurally identical, with the same smooth terms and the same random effects structure applied to each of the annual subsets.

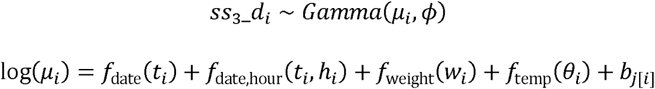

where:

μ_i_ is the expected swimming speed for observation i. f_date_(t_i_) is a smooth function of date (t_i_).

f_date,hour_(t_i_,h_i_) is a tensor product smooth of date (t_i_) and hour (h_i_), with thin-plate (tp) basis for date and cyclic cubic (cc) basis for hour.

f_weight_(w_i_) is a smooth function of weight (w_i_).

f_temp_(θ_i_) is a smooth function of mean temperature (θ_i_).

b_j[i]_∼N(0,σ^2^) represents a random intercept for fish j corresponding to observation i.

= is the shape parameter of the Gamma distribution.

#### 2.6.2 Depth use

The average daily depth of each individual fish was categorized into an ordered categorical variable with four levels: 0–0.5 m, 0.5–1 m, 1–1.5 m, and 1.5–4 m. This discretization was chosen to balance the design and avoid sparsely populated categories, as the deepest half-meter of the water column was rarely used in these ponds. To analyze depth utilization, we applied proportional-odds logit models, specifically cumulative link mixed-effects models (CLMMs) with a logit link function, to assess the effects of seasonality, time of day, and fish size on depth use across years. The models were fitted using the clmm() function from the R package ordinal (Christensen, 2019). We evaluated models with and without random intercepts for fish identity (fishID) to account for individual-level variability, as well as models with and without interactions among the covariates "month," "year," and "hour." Initially, a null model without covariates was compared to models with random effects to assess the importance of inter-individual variability. Subsequently, a series of models were fitted, incorporating combinations of the categorical covariates month (May–November), year (2022, 2023), and the continuous covariate hour. The best-supported model was determined based on Akaike’s Information Criterion corrected for small sample size (AICc). To test the influence of random effects for individual fish and interaction terms, likelihood ratio tests (LRTs) were used to compare nested models. Model fit was assessed using the McFadden pseudo R² and the selected model was used to estimate the probability of fish occupying specific depth ranges, with cumulative probabilities calculated based on odds ratios (ORs) between the levels of the dependent variables representing the relative likelihood of fish utilising different depths under different seasonal, temporal and interannual conditions as follows:

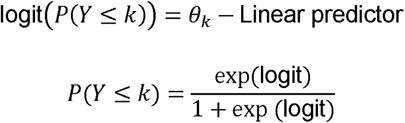

For each depth category, the probability was calculated as follows:

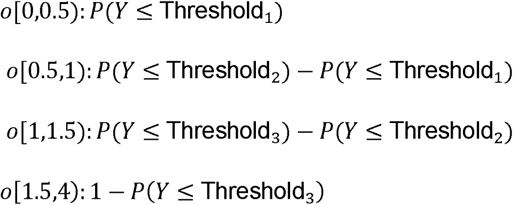

For further details on the parameterization of the CLMMs and the calculation and interpretation of the OR, we refer to other of our studies that follow a similar approach, (Říha et al., 2022).

#### 2.6.3 Feeding ground use

We analyzed feeding ground use (FGU) using Generalized Additive Models for Location, Scale, and Shape (GAMLSS) (Stasinopoulos and Rigby, 2007). FGU represents proportions ranging between 0 and less than 1, capturing the probability of foraging space use, with an FGU value of 0 indicating a passive foraging condition at the site. We used a zero-inflated beta distribution in the models to address the zero-inflation of this variable. Separate models were fitted for each year, undergoing model selection until convergence was achieved for one or more distribution parameters: μ, representing the mean of the distribution; σ, representing the scale or variance; and ν, representing the shape parameter, restricted in this case to account for the point mass at 0. Temporal trends in FGU were modeled by including a penalized P-spline smoothing function for the daily time variable. We also incorporated fish weight and the number of days since feeding as covariates, each adjusted using penalized P-splines, to account for the effects of size and feeding timing on the probability of foraging site use. Individual fish identity was modeled using a random smooth term to capture inter-individual variability. To control for residual autocorrelation in the time series, we included an autocorrelation structure in one or more submodels associated with the distribution parameters. Model selection, based on the Schwarz Bayesian Criterion (SBC), determined the final inclusion of these structures and smooth terms in the three potential distribution parameters (μ, σ, and ν), an approach allowed for flexibility and ensured the best-fitting model for each year. The models were implemented using the gamlss() function from the R package gamlss (Rigby and Stasinopoulos, 2009). Details on the selection process of the GAMLSS models and additional analyses can be found in the supplementary material, and further details on their parameterization can be found in (Říha et al., 2021).

#### 2.6.4 Food source utilization

To quantify the contribution of different food sources to the isotopic signatures of consumers, a separate Bayesian mixing model (Moore and Semmens, 2008) was run for each pond using the MixSIAR package (Stock and Semmens, 2016). The model for Loužek included four sources: zooplankton, macrozoobenthos, TF food sources, and cereals. Following the recommendations of (Zanden and Rasmussen, 2001), fractionation factors of 3.23 ± 0.41 ‰ for δ15N and 0.47 ± 1.23 ‰ for δ13C were used for animals, and 2.4 ± 0.42 ‰ for δ15N and 0.40 ± 0.28 ‰ for detritus and macrophytes (Mccutchan et al., 2003).

Diet composition was analyzed by calculating the median proportions of diet items (zoobenthos, TF sources, cereals, and zooplankton) consumed by carp in each year. To test for differences in the proportions of diet items between 2022 and 2023, we used the Wilcoxon rank-sum test, a non-parametric method suitable for comparing distributions between two independent groups.

#### 2.6.5 Growth evaluation

The effect of individual behavior on growth was tested using weight gain (i.e., the difference between stocking and harvest weight) as the response variable. Predictor variables included initial weight, mean feeding ground utilization (FGU; averaged during the feeding period - from beginning of feeding till one week after last feeding as a proxy for individual feeding ground use), mean swimming speed (as a proxy for individual activity), and year. Initial weight was included to account for potential size-dependent growth differences, while mean FGU and swimming distance were used to capture variations in feeding behavior and activity levels.

Additionally, growth and stable isotope data were available for more individuals (including those with malfunctioning tags). To make use of this broader dataset, a second model was constructed with growth as the response variable, and initial weight, proportion of supplementary grain consumption, and year as predictors. This approach allowed us to assess growth effects using a larger sample size, providing a broader understanding of growth patterns despite the incomplete telemetry data.

General Linear Models (GLMs) were used to assess the effect of each variable on growth. Model selection was performed using AIC to balance model complexity and goodness of fit, while ANOVA was used to test for significant effects among predictor variables, ensuring the most parsimonious and informative model was chosen.

All analyses were performed using R software version 4.2.0 (R Development Core Team, 2024).

## 3. Results

### 3.1 Swimming activity and utilization of depth and space

#### 3.1.1 Swimming activity

The final models for 2022 and 2023 shared similar predictors, including date, interaction between date and hour, mean temperature, and a random intercept for individual fish (Table 2, 1S). After release, swimming activity levels were similar between the two years, with an estimated mean of 0.036 m/s (95% CI: 0.025–0.046) in 2022 and 0.033 m/s (95% CI: 0.026– 0.042) in 2023. Seasonal activity patterns followed a consistent trend of steady decline across both years. Key differences emerged during July and August, with higher swimming activity observed in 2022 compared to 2023 (Fig. 2a).

**Figure 2.**
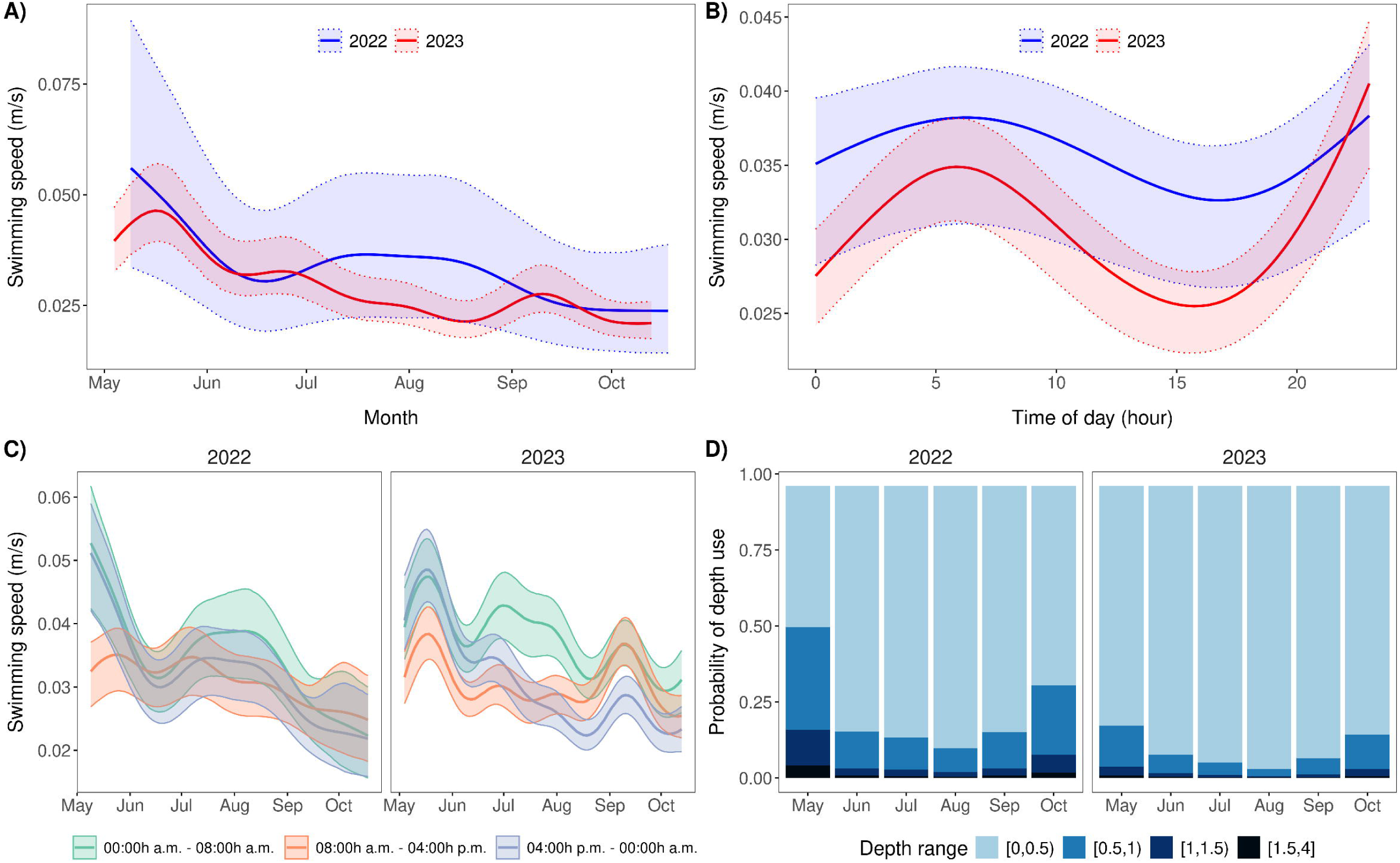
Temporal effects on fish activity (A, B, C) as estimated from Bayesian GAM models and estimates of depth utilization (D) from CLMM models in 2022 and 2023. (A) Seasonal trends in swimming speed across months. (B) Effect of time of day on swimming speed. (C) Monthly trend variations by time of day. Shaded areas represent 95% credible intervals. (D) Predicted probabilities of depth utilization over time (2022–2023) in the sampled ponds. The x-axis shows the seasonal trend in months (May to October), while the y-axis represents the probability of depth utilization. The depth ranges are differentiated in the legend: [0,0,5] (light blue) to [1,5,4] (dark blue). The probabilities were derived from the best-supported cumulative mixed link model (CLMM), which included fixed effects for year, month, hour and their interactions as well as random effects for individual fish. The highest probabilities for the use of deeper depths occurred in May and October, while shallower depths were more frequently used during the warmer summer months (June-August).

**Table 2.** Results of Bayesian additive mixed models analyzing swimming speed of carp in 2022 and 2023. Models were fitted separately for each year, incorporating smooth terms for temporal effects (date), environmental effects (temperature) and individual variability (fish weight), as well as a tensor product interaction between date and time of day (hour) to capture diel patterns of swimming activity. Random intercepts for individual fish (fishID) accounted for inter-individual variability in baseline swimming speed. The table summarizes the regression coefficients for parametric terms (total effects of covariates, with the intercept representing the log-transformed baseline swim speed), smoothing spline hyperparameters to quantify variability and smoothness of non-linear terms, and multilevel hyperparameters to represent inter-individual variability.

| Parameter | 2022 |  |  | 2023 |  |  |
| --- | --- | --- | --- | --- | --- | --- |
|  | Estimate | SE | 95% CI | Estimate | SE | 95% CI |
| **Parametric terms** |  |  |  |  |  |  |
| Intercept | -3.31 | 0.14 | [-3.68, -3.08] | -3.40 | 0.11 | [-3.63, -3.17] |
| **Smooth terms** |  |  |  |  |  |  |
| s(date) | -43.50 | 433.34 | [-851.37, 638.28] | -147.68 | 108.19 | [-316.30, 48.54] |
| t2(date × hour<sub>[00:00 a.m. - 06:00 a.m.]</sub>) | -0.05 | 0.04 | [-0.14, 0.02] | -0.02 | 0.04 | [-0.09, 0.06] |
| t2(date × hour<sub>[06:00 a.m. - 12:00 p.m.]</sub>) | 6.59 | 67.16 | [-99.08, 131.91] | 23.54 | 16.89 | [-7.03, 49.91] |
| t2(date × hour<sub>[12:00 p.m. - 6:00 p.m.]</sub>) | -0.03 | 0.02 | [-0.08, 0.02] | -0.09 | 0.02 | [-0.14, -0.06] |
| s(temp) | -1.70 | 0.98 | [-3.90, -0.06] | 0.34 | 0.53 | [-0.63, 1.43] |
| **Multilevel hyperparameters** |  |  |  |  |  |  |
| sd(Intercept) | 0.37 | 0.10 | [0.25, 0.68] | 0.53 | 0.10 | [0.38, 0.75] |
| **Smooth hyperparameters** |  |  |  |  |  |  |
| sds(date) | 1.29 | 0.53 | [0.52, 2.31] | 3.85 | 0.98 | [2.55, 6.37] |
| sds(t2(date × hour<sub>[00:00 a.m. - 06:00 a.m.]</sub>)) | 4.74 | 0.81 | [3.20, 6.36] | 7.71 | 1.75 | [3.94, 10.97] |
| sds(t2(date × hour<sub>[06:00 a.m. - 12:00 p.m.]</sub>)) | 4.88 | 1.40 | [2.78, 8.21] | 4.19 | 1.35 | [2.52, 6.88] |
| sds(t2(date × hour<sub>[12:00 p.m. - 6:00 p.m.]</sub>)) | 1.92 | 1.17 | [0.25, 4.97] | 2.70 | 1.07 | [0.92, 5.28] |
| sds(temp) | 1.16 | 0.34 | [0.58, 1.88] | 1.52 | 0.23 | [1.18, 2.09] |

In both years, daytime activity followed a similar pattern, with peaks occurring during the early night (around midnight) and early morning (around 5 a.m.). Activity declined in the late afternoon, reaching its lowest levels between 3 and 5 p.m. (Fig. 2B). However, the daytime activity pattern in 2022 was more consistent, with smaller fluctuations between high and low activity periods compared to 2023 (Fig. 2B, C). In 2022, there was moderate inter-individual variability in swimming activity, with an estimated standard deviation of baseline activity between individuals of 0.37 (SE = 0.10). This variability increased in 2023, reaching a standard deviation of 0.53 (SE = 0.10), suggesting greater behavioral diversity among fish in 2023.

#### 3.1.2 Depth utilization

Seasonal patterns in depth utilization were observed, with a strong preference for shallower depths in the warmer months. In June (difference from May=−1.62, p<0.001) and July (difference from May=−1.64, p<0.001), for example, the fish used significantly shallower depths. Conversely, in spring (May) and autumn (October), carp preferred greater depths, indicating seasonal shifts in habitat use (all p<0.01) (Table 3). Although there were no significant overall differences in depth use between 2022 and 2023 (contrast estimate = 1.04, SE = 0.809, p = 0.200), fish showed a general tendency to use shallower depths in 2023 (least square means (± SE): LSM_2023_ = -4.08 ± 0.51) compared to 2022 (LSM_2022_ = - 3.04 ± 0.63; Fig. 3). Cumulative probabilities derived from the model further supported these results, with fish showing a stronger preference for greater depths in 2022 (P(Yj≥3) = 0.012), corresponding to depths between 1 and 1.5 meters, than in 2023 (P(Yj≥3) = 0.004).

**Figure 3.**
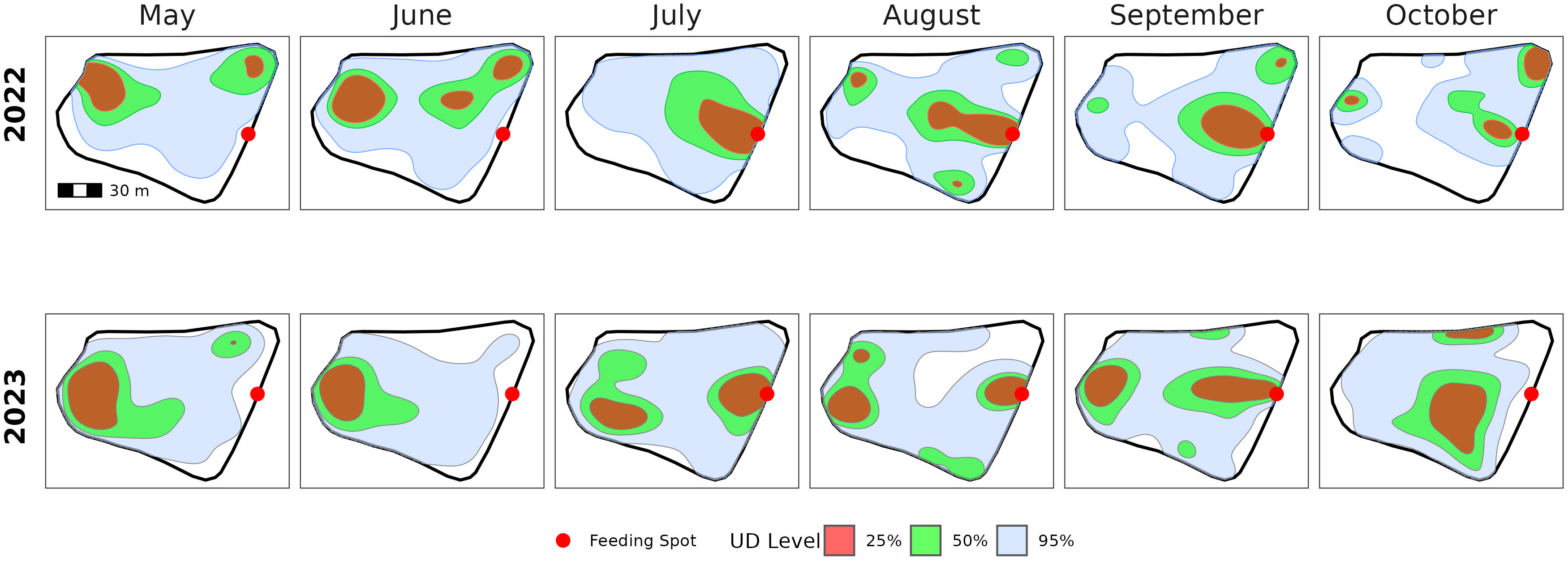
Spatiotemporal visualization of the activity centers of all tracked individuals (population kernel density estimation) across months for 2022 and 2023. The kernel density estimates represent utilization distributions (UD) of tracked individuals, with gray indicating 25% UD, green 50% UD, and blue 95% UD. The red dot marks the designated feeding ground. Each panel represents a month, with the top row displaying data for 2022 and the bottom row for 2023.

**Table 3.** Summary of cumulative link mixed model (CLMM) predicting carp use of depth over months and hours in 2022 and 2023. Depth is an ordinal dependent variable with thresholds corresponding to depth ranges, where θk is the cumulative probability of transition between depth categories. Fixed effects include year, month, hour and their interactions, where β1– β3 are the estimated regression coefficients for these variables. Random effects include individual fish identity (fishID) as a random intercept and residual variance (σ²). The intraclass correlation coefficient (ICC) reflects the degree of repeatability at the individual level. Marginal R² and conditional R² represent the variance explained by fixed effects alone or by fixed and random effects. * p < 0.05; ** p < 0.01.

| Variable | Estimate | S.E. | z-value | p-value |
| --- | --- | --- | --- | --- |
| **depth (thresholds)** |  |  |  |  |
| $\theta_1$ [0,0.5)\|[0.5,1) | -0.585 | 0.632 | -0.926 | 0.354 |
| $\theta_2$ [0.5,1)\|[1,1.5) | 1.136 | 0.632 | 1.798 | 0.072 |
| $\theta_3$ [1,1.5)\|[1.5,4] | 2.613 | 0.632 | 4.134 | <0.001 ** |
| **Year** |  |  |  |  |
| $\beta_1$ (2022, ref.) | - | - | - | - |
| $\beta_1$ (2023) | -1.625 | 0.811 | -2.003 | 0.045 ** |
| **Month** |  |  |  |  |
| $\beta_2$ (May, ref.) | - | - | - | - |
| $\beta_2$ (June) | -1.620 | 0.080 | -20.232 | <0.001 ** |
| $\beta_2$ (July) | -1.637 | 0.081 | -20.258 | <0.001 ** |
| $\beta_2$ (August) | -2.164 | 0.092 | -23.395 | <0.001 ** |
| $\beta_2$ (September) | -1.458 | 0.094 | -15.507 | <0.001 ** |
| $\beta_2$ (October) | -0.730 | 0.125 | -5.851 | <0.001 ** |
| **Hour** |  |  |  |  |
| $\beta_3$ | -0.045 | 0.004 | -10.969 | <0.001 ** |
| **Year × Month** |  |  |  |  |
| (2023 × June) | 1.014 | 0.111 | 9.127 | <0.001 ** |
| (2023 × July) | 0.764 | 0.115 | 6.659 | <0.001 ** |
| (2023 × August) | 0.125 | 0.134 | 0.934 | 0.351 |
| (2023 × September) | 0.688 | 0.124 | 5.571 | <0.001 ** |
| (2023 × October) | 0.743 | 0.153 | 4.845 | <0.001 ** |
| **Year × Hour** |  |  |  |  |
| (2023 × Hour) | 0.010 | 0.005 | 1.814 | 0.070 |
| **Month × Hour** |  |  |  |  |
| (June × Hour) | -0.015 | 0.006 | -2.497 | 0.013 ** |
| (July × Hour) | -0.032 | 0.006 | -5.178 | <0.001 ** |
| (August × Hour) | -0.012 | 0.007 | -1.762 | 0.078 |
| (September × Hour) | -0.035 | 0.007 | -5.020 | <0.001 ** |
| (October × Hour) | -0.012 | 0.009 | -1.285 | 0.199 |
| **Year × Month × Hour** |  |  |  |  |
| (2023 × June × Hour) | -0.020 | 0.009 | -2.349 | 0.019 ** |
| (2023 × July × Hour) | -0.026 | 0.009 | -2.814 | 0.005 ** |
| (2023 × August × Hour) | 0.022 | 0.010 | 2.189 | 0.029 ** |
| (2023 × September × Hour) | -0.004 | 0.010 | -0.454 | 0.650 |
| (2023 × October × Hour) | -0.013 | 0.011 | -1.173 | 0.241 |
| **Random Effects** |  |  |  |  |
| $\sigma^2$ | 3.913 | | | |
| $\tau_{00}$ (fishID) | 1.978 | | | |
| ICC | 0.34 |  |  |  |
| **Marginal R<sup>2</sup>/Cond. R<sup>2</sup>** | 0.25 / 0.54 |  |  |  |

Significant interactions between individual years and months also indicated different depth preferences in 2023 compared to 2022, particularly in June (estimate = 1.014, p < 0.001) and July (estimate = 0.764, p < 0.001). Fish depth use showed relatively low intraindividual variability, which is reflected in a high intraclass correlation coefficient (ICC = 0.796).

#### 3.1.3 Horizontal space use

The spatial patterns of horizontal space use were similar across both years (Fig. 3). Before the feeding period (May and June), the tracked individuals’ centers of activity (UD 50% or 25%) were primarily located in the western and northeastern parts of the pond, excluding the feeding ground. During the feeding period (July to September), the feeding ground became one of the key activity centers. After the feeding period ended (mid-September), the feeding ground was either partially included in the activity centers (2022) or fully excluded (2023). This trend was consistent across all tracked individuals in both years (Figs. 1 and 2 in Supplementary Material 2).

### 3.2 Utilization of feeding ground

Overall, the carp used the feeding ground more intensively in 2022 (FGU = 0.079) than in 2023 (FGU = 0.055), based on the predictions derived by the model from aggregated data. A significant time effect was observed in 2022 (b = -0.004, SE = 0.002, t = -2.06, p = 0.04) but not in 2023 (p > 0.1; Table 4). Feeding ground use increased early after the start of feeding (July-August), followed by a decline in the middle of the season (August-September), and recovered slightly towards the end of the season (September-October). For 2023, no significant temporal trend was observed (p = 0.78; Table 4), indicating a relatively stable FGU over the season with smaller fluctuations, compared to 2022 (Fig. 4A).

**Figure 4.**
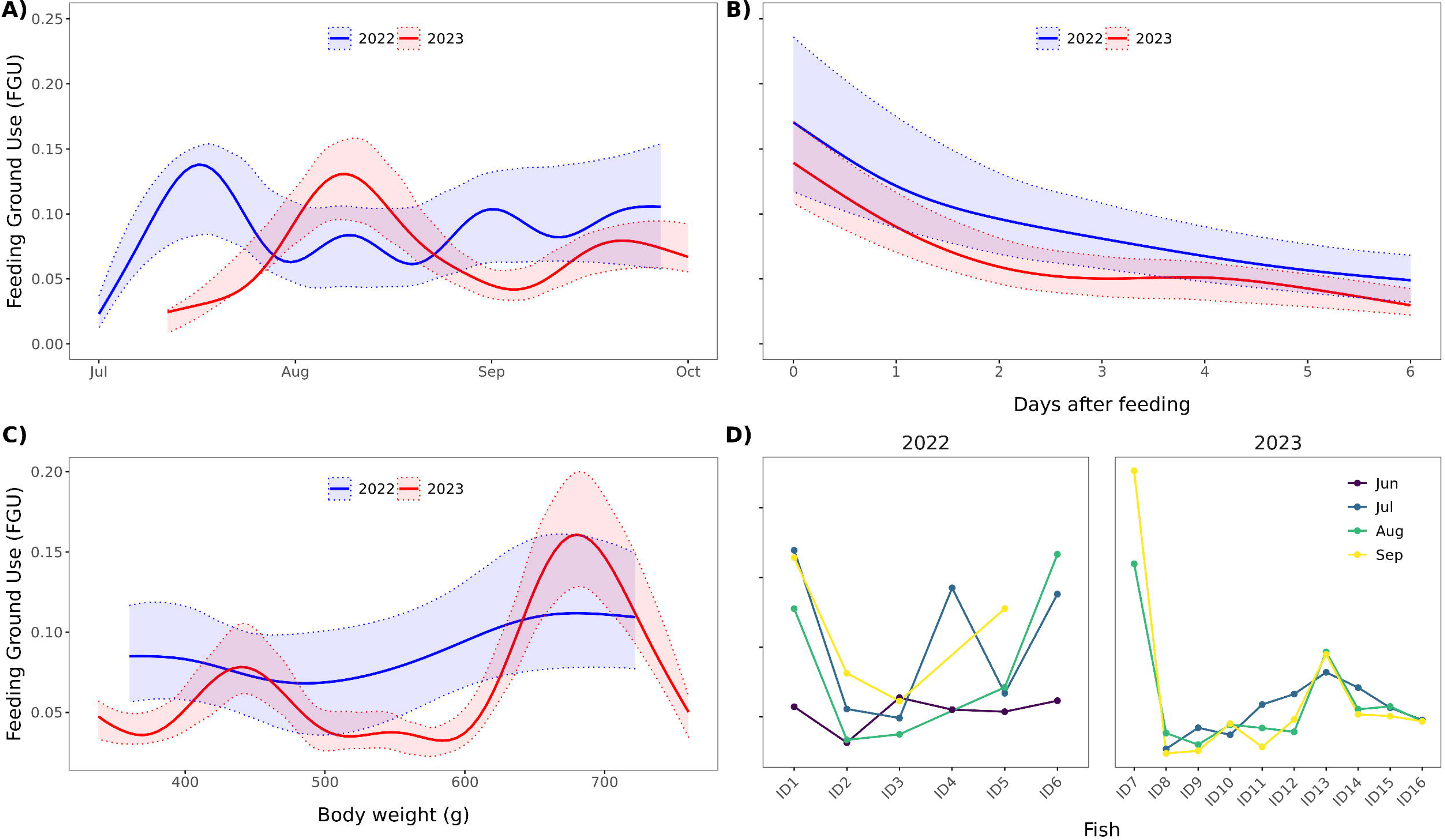
Seasonal patterns and relationships of feeding ground use (FGU) across 2022 (blue) and 2023 (red). (A) Seasonal trends in feeding ground use from July to October, with shaded areas representing 95% confidence intervals. (B) Decline in feeding ground use over the days following feeding events, showing a faster decrease in 2023 compared to 2022. The day of feeding is 0 and the day begins at the time of the actual feeding. (C) Relationship between body weight (g) and feeding ground use, with distinct trends observed for each year. (D) Individual variability in feeding ground use (FGU) by month, shown separately for 2022 and 2023. All estimates are based on bootstrap predictions from the fitted GAMLSS models for each year, accounting for covariates and random effects.

**Table 4.**
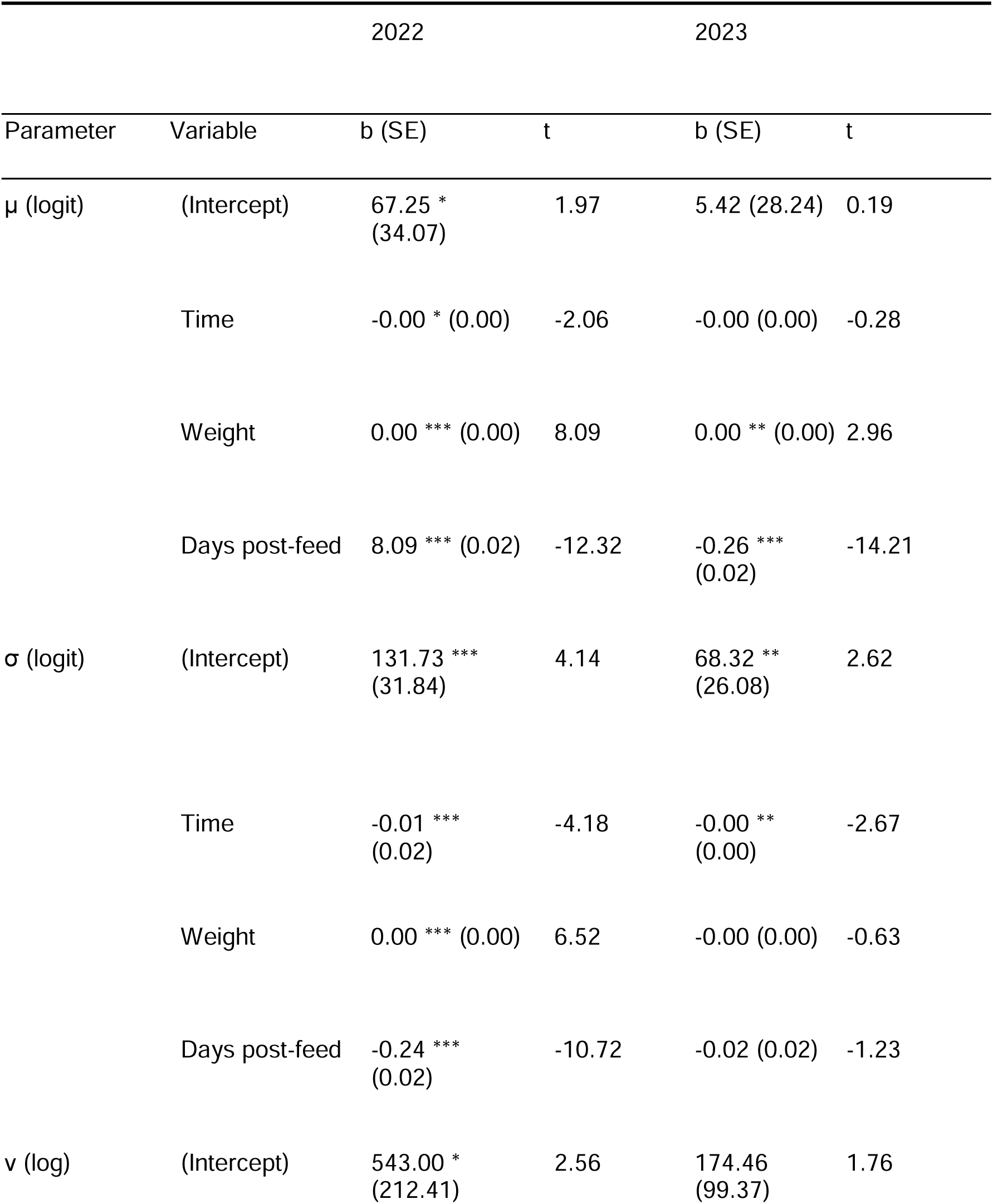
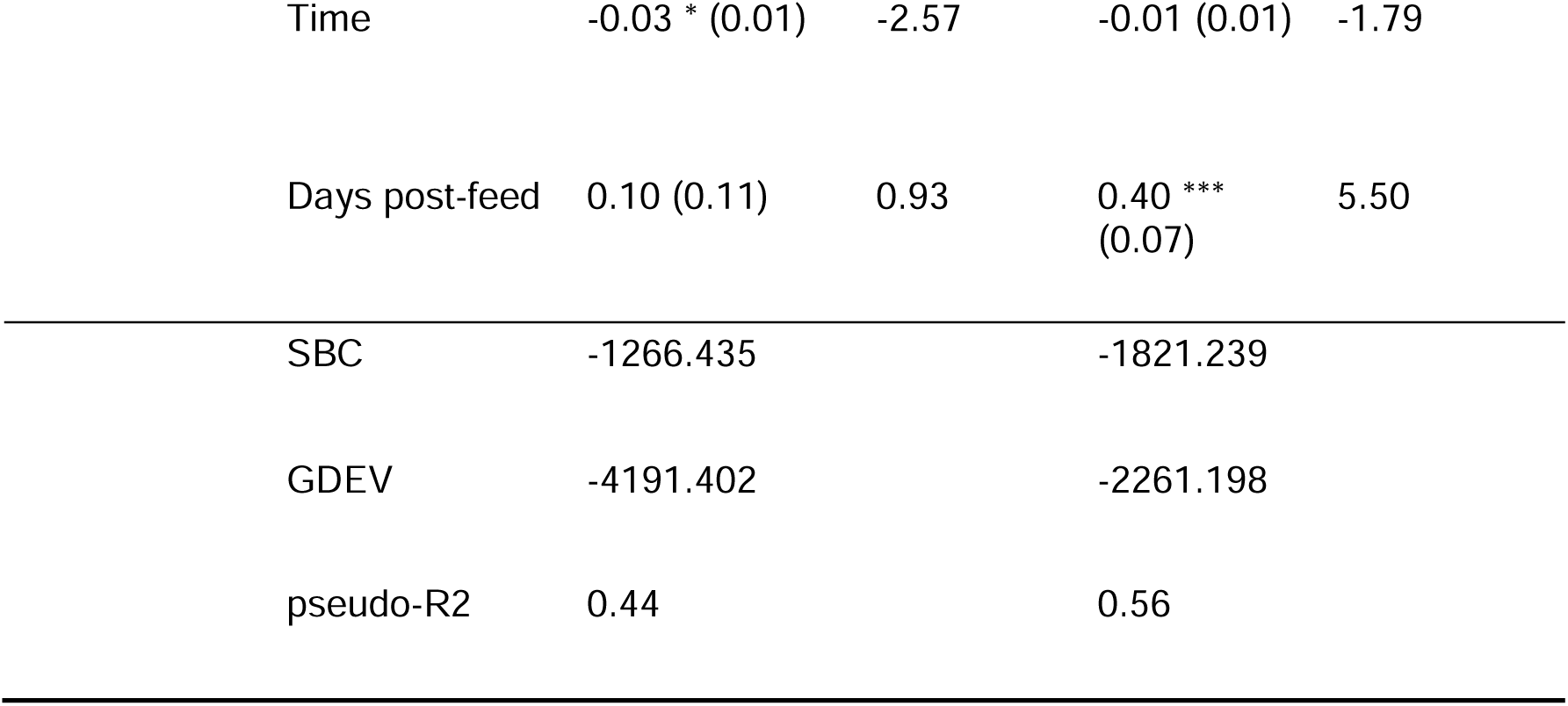
Summary of the zero-inflated beta GAMLSS model BEINF0 ∼ μ, σ, ν) analysing the effects of time, weight, and days after feeding on feeding ground use (FGU) in carp. Penalized P-spline (pb) smoothing terms were used to model covariates, and random effects were incorporated with a random intercept and autocorrelation structure across all three distribution parameters (μ,σ,ν). b, variable estimate (z-score); SE, standard error; t, t-statistic; edf, effective degrees of freedom of smooth terms; GDEV, global deviance; SBC, Schwartz Bayesian Criterion; pseudo-R^2^; * p < 0.05; ** p < 0.01.

|  |  | 2022 |  | 2023 |  |
| --- | --- | --- | --- | --- | --- |
| Parameter | Variable | b (SE) | t | b (SE) | t |
| $\mu$ (logit) | (Intercept) | 67.25 *<br>(34.07) | 1.97 | 5.42 (28.24) | 0.19 |
|  | Time | -0.00 * (0.00) | -2.06 | -0.00 (0.00) | -0.28 |
|  | Weight | 0.00 *** (0.00) | 8.09 | 0.00 ** (0.00) | 2.96 |
|  | Days post-feed | 8.09 *** (0.02) | -12.32 | -0.26 ***<br>(0.02) | -14.21 |
| $\sigma$ (logit) | (Intercept) | 131.73 ***<br>(31.84) | 4.14 | 68.32 **<br>(26.08) | 2.62 |
|  | Time | -0.01 ***<br>(0.02) | -4.18 | -0.00 **<br>(0.00) | -2.67 |
|  | Weight | 0.00 *** (0.00) | 6.52 | -0.00 (0.00) | -0.63 |
|  | Days post-feed | -0.24 ***<br>(0.02) | -10.72 | -0.02 (0.02) | -1.23 |
| $\nu$ (log) | (Intercept) | 543.00 *<br>(212.41) | 2.56 | 174.46<br>(99.37) | 1.76 |

|  |  |  |  |  |
| --- | --- | --- | --- | --- |
| Time | -0.03 * (0.01) | -2.57 | -0.01 (0.01) | -1.79 |
| Days post-feed | 0.10 (0.11) | 0.93 | 0.40 ***<br>(0.07) | 5.50 |
| SBC | -1266.435 |  | -1821.239 |  |
| GDEV | -4191.402 |  | -2261.198 |  |
| pseudo-R2 | 0.44 |  | 0.56 |  |

In both years, the proportion of FGU (μ; logit scale) was significantly influenced by days after feeding and body weight (Table 4). Days after feeding had a significant negative effect in 2022 (b = -0.31, SE = 0.02, t = -12.32, p < 0.001) and 2023 (b = -0.26, SE = 0.02, t = - 14.21, p < 0.001), indicating that FGU declined sharply during the first two days after feeding, after which the rate of decrease slowed (Fig. 4B). Body weight had a positive influence on FGU in both years (2022: b = 0.0029, SE = 0.0004, t = 8.09, p < 0.001; 2023: b = 0.001, SE = 0.0002, t = 2.96, p < 0.01; Fig. 4C) indicating bigger carps utilized the feeding ground more intensively.Variability in FGU (σ; logit scale) was also significant in both years (Table 4). In 2022, FGU variability had strong negative correlations with time (b = -0.007, SE = 0.002, t = -4.18, p < 0.001) and days after feeding (b = -0.24, SE = 0.02, t = -10.72, p < 0.001). In 2023, variability was significantly influenced by time (b = -0.004, SE = 0.001, t = - 2.67, p < 0.01), but neither weight (p = 0.53) nor days after feeding (p = 0.22) contributed significantly. The probability of zero utilization of the feeding ground (ν; logarithmic scale) showed a significant negative effect of time in 2022 (b = -0.03, SE = 0.01, t = -2.57, p < 0.05), while days after feeding had no significant impact (p = 0.35). In 2023, days after feeding had a strong positive effect (b = 0.40, SE = 0.07, t = 5.50, p < 0.001), while temporal effects were only marginally significant (b = -0.009, SE = 0.005, t = -1.79, p = 0.07).

Inter-individual variation in FGU was substantial and similar in both years, with individual average FGU (over the season) ranging from 0.05 to 0.14 in 2022 and 0.03 to 0.18 in 2023 (Fig. 4D). The extent of inter-individual variation was significantly greater in 2022 (t-test: t = 4.07, p = 0.001) than in 2023 (among monthly averages 22.9-62.4% and 1.1-35.3% in 2022 and 2023, respectively; Fig. 4D).

### 3.4 Natural food availability

The biomass of benthic invertebrates followed a similar seasonal trend in both years, with the highest densities observed in June. This was followed by a sharp decline from July to early September (to values close to zero in 2022) and a slight elevation by the end of September (Fig. 5A).

**Figure 5.**
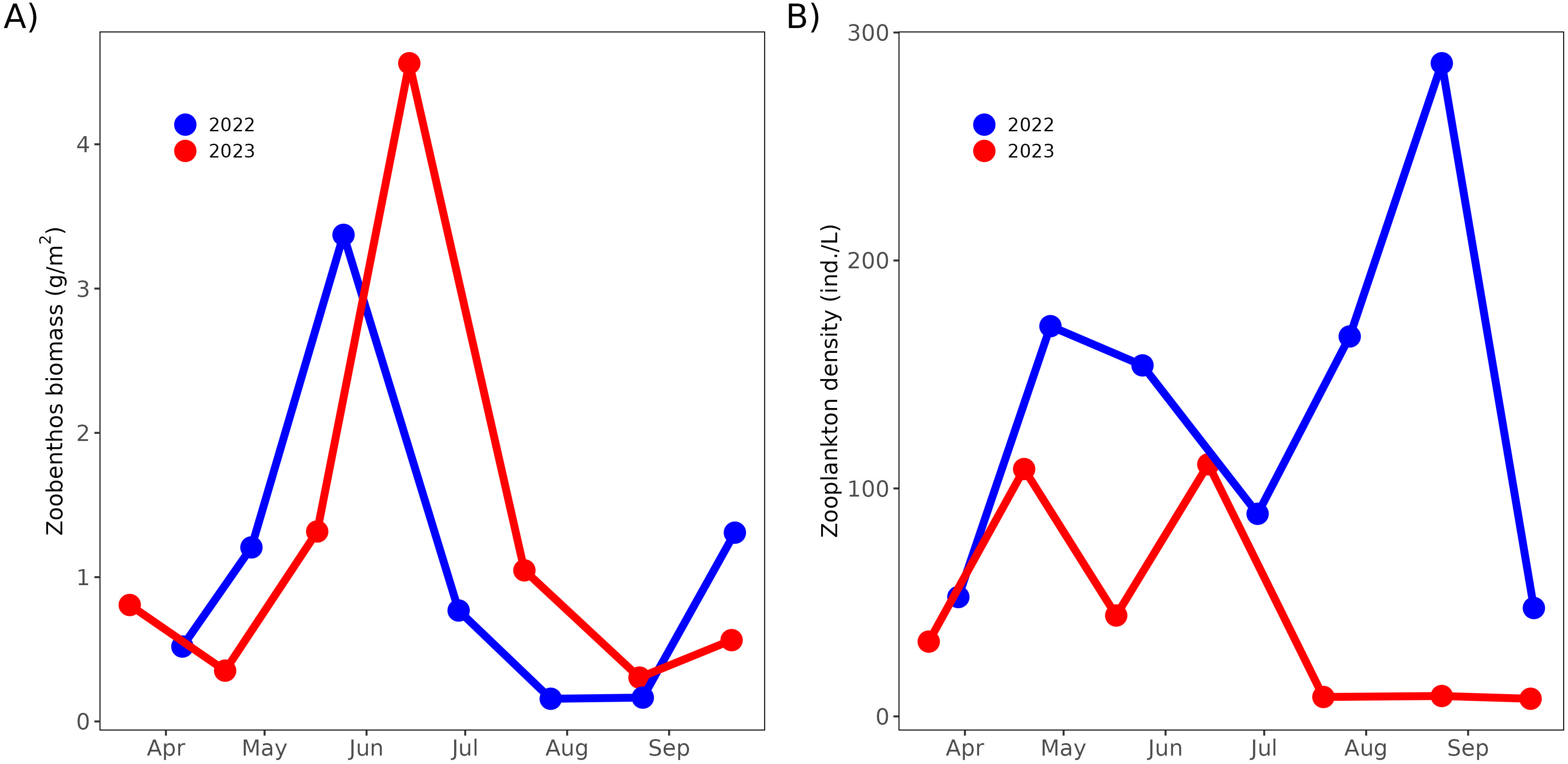
Seasonal trends in zoobenthos dry weight (A) and densities of zooplankton larger than 200 µm (B). Panel A shows the zoobenthos dry weight (g/m²), while panel B illustrates zooplankton density (individuals/L) for total zooplankton.

The seasonal trends in the development of the zooplankton (>200 µm) abundance varied from year to year (Fig. 5B). Until July, the trend in densities was relatively similar. After July 2023, however, the zooplankton density decreased to almost zero. In contrast, zooplankton density in 2022 increased in August and decreased in September (Fig. 5B).

### 3.5 Diet and growth of tagged individuals

#### 3.5.1 Diet

The dominance of diet items in carp was broadly consistent across both years. Zoobenthos had the highest median proportion (2022: 0.48, 2023: 0.49), followed by TF sources (2022: 0.20, 2023: 0.33), followed by cereals (2022: 0.18, 2023: 0.15), and zooplankton (2022: 0.14, 2023: 0.03; Fig. 6). Statistically significant differences between years were found for the share of zooplankton (*p* < 0.001) being higher in 2022 (*p* = 0.0499), and TF sources in 2023. No significant inte-year differences were observed for zoobenthos or cereals.

**Figure 6.**
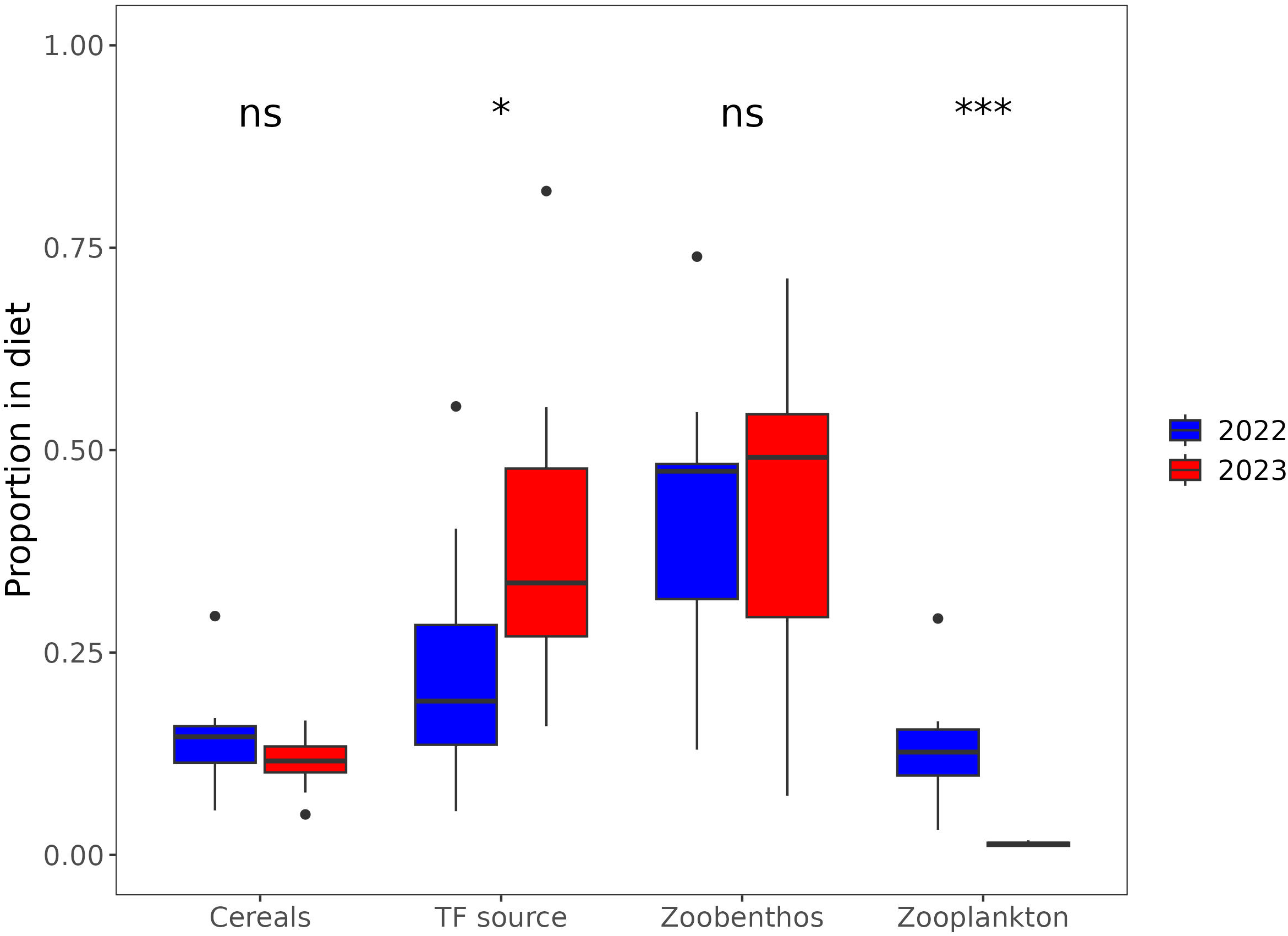
Contribution of different food sources to the diet of recovered tagged carps in 2022 (blue) and 2023 (red). The boxplots represent the proportion of each food source in the diet: benthos, cereals, TF sources, and zooplankton. Statistical significance of differences between years is indicated above the categories: ns (not significant), * (p < 0.05), and *** (p < 0.001).

#### 3.5.2 Growth

The most parsimonious model indicated that both year and Feeding Ground Use (FGU) significantly influenced individual growth (weight gain) based on individuals with available telemetry data. Growth was significantly higher in 2023 (mean weight gain: 1071 ± 219 g) compared to 2022 (617 ± 179 g; *p* < 0.001; Fig. 7A). Additionally, growth showed a significant positive relationship with FGU (*p* = 0.015; Fig. 7B).

**Figure 7.**
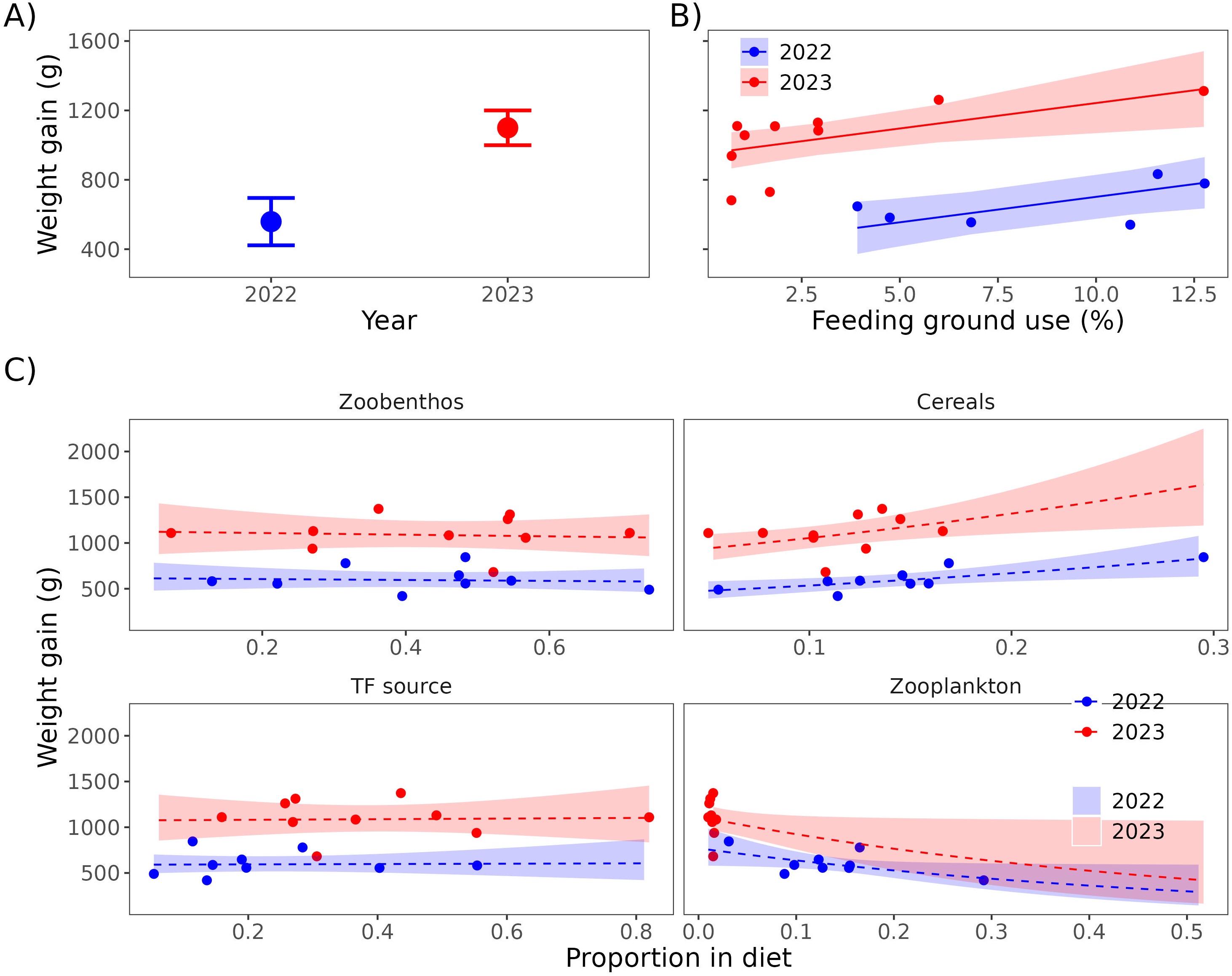
Tagged individuals weight gain and its relationship with feeding ground use and dietary composition for 2022 and 2023. (A) Average weight gain (g) with standard errors for each year. (B) Relationship between feeding ground use (FGU; %) and weight gain (g), with shaded areas representing 95% confidence intervals. (C) Relationships between dietary proportions of benthos, cereals, TF sources and zooplankton and weight gain (g), showing separate trends for 2022 (blue) and 2023 (red) with 95% confidence intervals.

Models evaluating growth with diet items consumed showed that growth differed significantly between years (higher in 2023) and correlated positively with the proportion of cereals (Table 5; Fig. 7C). The effect of cereal consumption on growth showed a consistent slope in both years, as the most parsimonious model was additive (Table 5). Other dietary components showed no significant correlation with growth (Table 5).

**Table 5.** Summary of GLM results evaluating the effects of dietary composition (proportions of cereals, benthos, zooplankton, and TF sources) and year (2023 compared to 2022) on the weight gain. For each diet variable, the table includes the intercept, estimated effect size (Estimate), standard error (S.E.), t-value, and p-value. Significant p-values are indicated as follows: *** (p < 0.001), ** (p < 0.01), * (p < 0.05), and . (p < 0.1).

| Diet | Variable | Estimate | S.E. | t-value | p-value |
| --- | --- | --- | --- | --- | --- |
| **Cereals** | Intercept | 6.0561 | 0.1368 | 44.271 | < 0.001 *** |
|  | Grain prop. | 2.2573 | 0.8404 | 2.686 | 0.016 * |
|  | Year (2023) | 0.6788 | 0.0859 | 7.905 | < 0.001 *** |
| **Benthos** | Intercept | 6.4227 | 0.1371 | 46.865 | < 0.001 *** |
|  | Bent. prop. | -0.0832 | 0.2788 | -0.299 | 0.769 |
|  | Year (2023) | 0.605 | 0.0976 | 6.197 | < 0.001 *** |
| **Zooplankton** | Intercept | 6.6463 | 0.1437 | 46.262 | < 0.001 *** |
|  | Zoo. prop. | -1.888 | 0.9408 | -2.007 | 0.062 . |
|  | Year (2023) | 0.3709 | 0.1454 | 2.55 | 0.021 * |
| **TF sources** | Intercept | 6.3805 | 0.0979 | 65.204 | < 0.001 *** |
|  | Ter. in | 0.0313 | 0.292 | 0.107 | 0.916 |
|  | Year (2023) | 0.599 | 0.1087 | 5.512 | < 0.001 *** |

### 3.6 Harvest of the stock

During the harvest in 2022, 714 carp were caught with an average weight of 845 g, an average weight gain of 652 ± 223 g and a total biomass of 603 kg. In 2023, 820 carp were harvested, with an average weight of 887 g, an average weight gain of 778 ± 204 g and a total biomass of 727 kg. In addition, 13,900 carp (mean weight: 48 g; total biomass: 670 kg, newcomers with the flood) and 32 kg of topmouth gudgeon (*Pseudorasbora parva*) were caught in 2022 and 400 kg of topmouth gudgeon in 2023. As a result, the total biomass of carp at the end of the 2022 season was 1.75 times higher than in 2023, while the total fish stock in 2022 was only 1.16 times higher.

## 4. Discussion

Our study shows that carp activity, depth utilisation and horizontal space utilisation exhibit different temporal patterns reflecting seasonal and diurnal cycles and feeding regimes. Furthermore, our results underline the role of interannual variability, with carp behaviour responding dynamically to specific environmental conditions. In terms of feeding ground utilisation, all individuals tracked visited the feeding ground to varying degrees, with utilisation strongly influenced by individual differences, annual variation and time elapsed since feeding. Our results suggest that the ability of individuals to use the feeding site increases the intake of supplementary food, which in turn increases individual growth rates. These findings highlight the high behavioural plasticity of carp, their ability to dynamically adapt to local conditions and the significant influence of individual traits on growth performance.

The testing of our study hypotheses was influenced by two significant issues that affected the expected outcomes. First, an unexpectedly high rate of tag malfunctions in 2022 resulted in a lower number of tracked individuals, which limited the sample size for that year. This limitation prevented a more robust estimate from models and introduced a certain degree of variability among individuals in key parameters, resulting in excessively wide confidence intervals, and underperformance of models. Despite these challenges, the 2022 data are still valuable as they provide evidence of important behavioral adaptations in carp in response to contrasting environmental conditions across years. It was facilitated by the application of a Bayesian framework that helped to deal with uncertainty more effectively than conventional techniques (Gelman et al., 2013). The most striking difference between 2022 and 2023 was the fish biomass in the pond. Following a flood event in June 2022, the carp biomass nearly doubled, drastically increasing competition for food resources, and inevitably influenced the whole study background from natural prey availability over amount of grain feeding available for target carp to (not only) tagged carp behaviour and growth. The extent of this June flood-originated situation was revealed just at the harvest of the pond in October.

### 4.1 Activity

Our results reveal a distinct seasonal pattern of activity, characterized by a decreasing trend over the course of the season in both years. Activity levels were initially similar immediately after stocking in both years but diverged during the summer, with elevated activity observed in 2022. This divergence diminished by autumn, with activity levels converging again. A similar pattern of high spring activity (end of April and beginning of June), followed by a slight reduction at the onset of summer, has been documented under natural conditions and is linked to spawning activity (Banet et al., 2022; Watkinson et al., 2021). However, the carps in our study had not yet reached maturity, and no spawning activity was observed. Consequently, the high post-stocking activity observed in this study may be associated with the acclimatization process of fish to their new environment. Comparable post-translocation activity increases have been reported in catfish (Monk et al., 2020).

The onset of elevated summer activity in 2022 coincided with a flood event that increased the carp stock in the pond (see above in chapter 4). The prolonged increase in activity observed during the summer of 2022 aligns with previously documented effects of food shortages on fish behavior (Orpwood et al., 2006; Říha et al., 2024; Sogard and Olla, 1996).

Our results indicate that this increase coincided with a lower availability of both natural and supplementary food sources during the same period (see below). In general, reduced food availability triggers increased individual activity as fish seek to balance resource scarcity (Orpwood et al., 2006; Říha et al., 2024; Sogard and Olla, 1996).

The decline in activity in autumn corresponds to a drop in temperatures, from 18°C in early September to 13°C in mid-October. This is consistent with the temperature-dependent nature of carp activity (Cooke and Schreer, 2003). A similar decline transitioning into low winter activity has already been documented in natural settings (Banet et al., 2022; Bauer and Schlott, 2004). In summary, our findings indicate that both regular seasonal temperature fluctuations and inter-annual variations in resource availability dynamically influence the seasonal activity patterns of carp. These findings highlight the importance of environmental factors, such as temperature and resource availability, in shaping fish activity at both intra-and inter-annual scales.

Our results revealed consistent diurnal activity patterns in both years, with carp exhibiting the highest activity during the early night and morning and the lowest activity during the late afternoon. These findings align with previous studies reporting peak activity during nocturnal and twilight periods (Bajer et al., 2010; Benito et al., 2015; Hundt et al., 2022; Žák, 2021). Comparative analysis between 2022 and 2023 indicated reduced diurnal fluctuations in activity (differences between minimal and maximal activity throughout the day) in 2022. This pattern corresponds to the increased overall activity levels observed during that year. Uniform diel activity patterns are commonly associated with resource scarcity or density-dependent behaviors, as documented in other fish species (Fingerle et al., 2016; Hansen and Closs, 2005; Říha et al., 2024). Our findings suggest that carp at higher densities may adjust both the rate and timing of their activity to compensate for limited food resources and optimize food acquisition. Such adjustments highlight behavioral plasticity that enables carp to adapt to resource constraints, potentially mitigating to some extent competition in higher-density settings.

### 4.2 Depth

Except for May and, to a certain extent, October, carp primarily utilized the very shallow areas of the pond (down to a depth of 0.5 meters) in both years of observation, with a stronger preference for these shallow zones in 2023. Information on depth use by carp in pond environments is sparse, and to our knowledge, no relevant studies have addressed this topic comprehensively. Even in natural environments, available data are very limited. Only Benito et al. (2015) studied a daily depth range used by carp in reservoirs across whole the year. The authors showed that carp favour a shallower part during the warm season and move deeper as the temperature decreases. However, the depth utilisation they observed has a very strong diurnal aspect, especially in summer and broader deoth range. Our findings suggest that the relatively uniform bathymetry of the pond, where depths exceeding 1 meter account for approximately 50% of the pond’s surface, results in large portions of the pond bottom being underutilized during summer. This behavior may be linked to oxygen availability, as oxygen depletion often occurs in deeper parts of ponds during summer. Oxygen concentration is a critical factor influencing fish habitat and depth use (Vejřík et al., 2016), even though carp is relatively tolerant of low oxygen conditions (Zhou et al., 2000) and it can even utilized hypoxic conditions (Benito et al., 2015). Unfortunately, due to technical issues, we could not obtain oxygen concentration profiles and therefore could not directly assess its effects on carp distribution. However, the pond was not hypertrophic and there was no strong algal bloom or oxygen supersaturation near the surface, which would have led to conditions causing profound oxygen depletion. Nevertheless, the possibility of localized oxygen limitations in deeper areas cannot be entirely excluded.

Another potential driver of frequent shallow water use can the regulation of body temperature or food availability. Nordahl et al. (2018) demonstrated that carp often engage in sunbathing near the surface, which allows them to raise their body temperature above ambient water temperatures, significantly enhancing their growth. In 2023, this behavior may have been especially beneficial given the higher prey availability, as elevated temperatures combined with improved nutrition could have maximized growth potential (see below in chapter 4.5). Beside that, the bottom in deeper parts of of ponds typically contain less macrozoobenthos (both biomass and density) compared to the littoral zones (Kajgrova et al., 2021). Shallow areas in and around the littoral zone, therefore, provide more abundant food resources than the pelagic zones (macrophyte-free areas) of ponds. In conclusion, while our findings indicate key behavioral trends and potential drivers of depth use by carp, more detailed studies are needed to fully understand the effects of ecosystem factors (e.g., oxygen, temperature, food distribution) and internal drivers (e.g., growth and metabolic demands).

### 4.3 Feeding ground utilization

Our results showed that the carp responded rapidly to supplementary feeding and actively utilized the feeding ground during the feeding period. This fast response aligns with previous research demonstrating that carp possess strong learning abilities and reliable spatial memory, enabling them to quickly identify and exploit food resource patches (Bajer et al., 2010; Hundt et al., 2022; Monk and Arlinghaus, 2017). Outside of the scheduled feeding times, however, the feeding ground remained unused. It is likely that conditions near the dam—such as steep shorelines and greater water depth—rendered the area less attractive compared to shallower regions closer to the tributary or in the northeast corner of the pond (Kajgrova et al., 2021). Moreover, each carp exhibited a relatively stable alternative activity area besides the feeding ground during the feeding period, a pattern observed consistently across all tracked individuals. This finding corresponds to our additional results, which showed that feeding ground use decreased relatively quickly over short time scales. In both study years, carp made intensive use of the feeding ground on the first day after feeding (on average 15% and 12% of their time in 2022 and 2023, respectively), followed by a gradual decline to about 5% and 3% of their time by the sixth day after feeding. Similar levels of time spent near feeding grounds were documented by Mehner et al. (2019), and a low use of feeding ground 4 days after feeding was noted by Jurajda et al. (2016). These results strongly suggest that all tracked carp individuals optimize their foraging activity in response to resource density. When supplementary feeding is quickly depleted, the carp return to their customary activity centers to forage for natural food sources. This adaptive foraging strategy underscores the importance of both spatial and temporal variability in resource availability, as well as the carp’s ability to learn and adjust their behavior accordingly.

### 4.4 Diet

Benthic invertebrates emerged as the most important dietary component for carp, followed by TF sources, cereals and zooplankton. The prominence of benthic macroinvertebrates in the diet of carp is well-documented in numerous studies (Anton-Pardo et al., 2014; Anton-Pardo and Adámek, 2015; García-Berthou, 2001; Rahman, 2015). However, the relatively high proportion of insects observed in this study is somewhat unexpected, as they typically constitute only a minor fraction of the carp diet (Lammens and Hoogenboezem, 1991). This anomaly may be explained by the favorable relationship of the pond size to its shoreline length in combination with the character of its surrounding landscape (farmland and trees).

In contrast, the relatively low contribution of zooplankton in the diet are consistent with previous research, which indicates a decline in zooplankton importance as carp undergo ontogenetic development as well as their depletion in ponds at the end of the season (Anton-Pardo and Adámek, 2015; García-Berthou, 2001; Rahman, 2015). In the size class studied (> 300 mm TL), zooplankton typically comprises only a minor portion of the diet, but this depends on food availability and spatial and temporal variation (Anton-Pardo et al., 2014; Anton-Pardo and Adámek, 2015; García-Berthou, 2001).

Further, our results showed that the zooplankton consumption is in a direct relation to the composition of the zooplnaton community present in the pond environment. The reduced zooplankton contribution observed in 2023 coincides with the near absence of larger zooplankton (> 200 µm) in the pond, emphasizing limited availability of this food resource. This pattern is likely linked to the high density of invasive topmouth gudgeon (*Pseudorasbora parva*). As an efficient zooplankton feeder, this small species can substantially deplete zooplankton populations and effectively outcompete carp in pond environments (Kajgrová et al., 2022; Musil et al., 2015).

Our results stress the crucial role of cereals in promoting the rapid growth of carp. While stable isotope analysis suggests that cereals constitute only a relatively minor proportion of the diet, their actual importance may be underestimated. This discrepancy arises because cereals primarily serve as an energy source rather than being utilized for protein synthesis or other anabolic processes, which are more strongly reflected in the nutrient composition of tissues and, consequently, in stable isotope signatures (Martínez Del Rio et al., 2009; Post, 2002). The significant positive relationship observed between the proportion of cereals in the diet and weight gain underscores their critical role as an energy source. Cereals, being rich in carbohydrates, provide an efficient and easily digestible source of energy for carps (∼2759.4 kcal.kg^−1^; Roy et al., 2020) having an energy profile near to their optimum requirement (∼3200 kcal.kg^−1^ diet; NRC, 2011). In contrast, other dietary components, such as benthic and TF sources or zooplankton, did not show such a similar relationship with growth. While cereals appear to drive growth through their energetic contribution, natural food sources such as zooplankton and benthic invertebrates likely play a complementary role. These protein-rich sources provide the essential building blocks (amino acids) required for tissue synthesis. Carp likely allocate the energy derived from cereals to meet metabolic demands, thereby sparing protein obtained from natural food sources for growth-related processes.

### 4.5 Growth

In general, the average individual growth was lower in 2022 than in 2023. This result seems to reflect the differences in stock density and respective food availability between years, as the stock density of carp was 1.75 times higher in 2022, while the level of co-feeding was similar. As a result, benthic food resources were depleted faster (almost no benthic food was available between July and September), resulting in lower per capita food availability in 2022.

Increased stock density can lead to several density-dependent effects that directly influence fish growth and behavior. High densities intensify competition for resources, which not only reduces per capita food availability but also increases the energetic costs associated with foraging and social interactions (Matte et al., 2021). In carp, aggressive interactions and competitive exclusion at feeding grounds have been shown to disproportionately affect subordinate individuals, leading to significant inequality in resource access and growth rates (Hundt et al., 2022; Jurajda et al., 2016).

Our results also suggest that the carp displayed more extensive and consistent activity in the summer of 2022, leading most likely to higher energy expenditure. Previous research suggests that increased activity can partially compensate for reduced food availability (Orpwood et al., 2006; Říha et al., 2021). However, in our case, the increased carp activity could not compensate for the limited food supply and may even have exacerbated the decline in growth (Hansen and Closs, 2005). Supplementary feed comprised only 11 g per 1 kg of carp per day during the feeding period in 2022 and 23 g per 1 kg of carp per day in 2023. Optimal feeding ranges between 2–5% of body weight per day (Hartman and Regenda, 2016), and sub-optimal feeding is not sufficient to cover the increased energy requirements during increased activity. A more precise calculation of the energy budget is necessary to fully understand the interplay between activity variation, food availability and growth. However, our results suggest that behavioural adaptations to fluctuating food resources can have a significant impact on production yield, especially under conditions of limited food availability.

Individual behavioural variations significantly influenced the carp growth. In both years, growth was positively related to the time spent at the feeding ground and the proportion of supplementary food consumed. Previous research has shown considerable individual heterogeneity and inequality in the feeding behaviour of carp with individuals spending more time at the feeding grounds tend to grow heavier (Hundt et al., 2022; Jurajda et al., 2016). Such a feature is likely to be related to carp personality with bolder carp being shown to monopolise spatially restricted feeding sites (Huntingford et al., 2010) and some individuals being excluded from access to the feeding ground at all (Jurajda et al., 2016). These findings could explain the patterns observed in our study, where certain individuals, potentially possessing higher competitive abilities, were able to stay longer at the feeding ground achieving better individual growth. Thus, individual traits, such as boldness, aggression, or exploratory behavior, can play an important role in determining access to resources and, consequently, growth rates.

### 4.6 Outcomes for aquaculture

Our results imply following important recommendations for carp aquaculture and that could contribute to more effective carp production:

Multiple feeding grounds -Even in small ponds with adequate spacing such a measure can help reduce competition among fish, ensuring that even less dominant individuals have access to food and minimize growth disparities.

Basic monitoring of natural food availability across the growing season -This can help to better plan feeding regime to maintain optimal feeding rate range of 2–5% of body weight per day and sufficient growth rate. However, one has to keep in mind that overfeeding using cereals is undesirable towards the end of the growing season both because of unwanted lipogenesis and increased negative nutrients load to the pond ecosystem. In our case, it would be advantageous to adjust the dose of supplementary feeding to reflect natural food availability and to increase it in August when the natural food resources are depleted. Providing protein-enriched food during this period could help to compensate for the lack of proteins and carbohydrates from natural sources and prevent the deterioration of fatty acid profiles in muscle tissue.

To reduce the density of invasive species like topmouth gudgeon and prevent their massive occurrence, as they can effectively compete with carp for natural prey.

## 5. Conclusions

Our study highlights the remarkable behavioural plasticity of carp and demonstrates their ability to adapt to different environmental conditions both within and between years. We found that carp activity, depth utilisation and feeding behaviour are shaped by seasonal and diurnal cycles, interannual variation and individual traits, all of which contribute to growth performance and aquaculture outcomes. The pronounced contrast in carp behaviour and growth between 2022 and 2023 illustrates the profound influence of complex interactions of food availability, stocking density and environmental conditions on production efficiency.

From an aquaculture perspective, our findings provide actionable insights to improve carp farming practices. Establishing multiple feeding sites, adapting feeding to natural food availability and optimising the distribution of supplementary feed during critical periods (e.g. late summer) can mitigate competition and promote growth. These measures could reduce the impact of density-dependent effects and ensure more equitable access to resources among individuals, leading to better overall yield and more sustainability.

Despite some methodological challenges, our study emphasises the crucial role of state-of-the-art research methods in advancing aquaculture science. High-resolution telemetry and stable isotope analysis provide invaluable insights into the interactions between the spatial and trophic behaviour of farmed animals and their environment. Equally important is the monitoring of other trophic levels utilised by the farmed animals, especially in semi-natural aquaculture systems that combine natural food resources with supplementary feeding. These methods can be further enriched by integrating novel tools such as genetic profiling, physiological assessments or microbiome analyses (Araujo et al., 2022; Brosset et al., 2021; Parata et al., 2021) that allow a comprehensive understanding of aquaculture dynamics. Integrating these advanced techniques not only allows for detailed descriptions of animal behavior and trophic interactions but also provides mechanistic insights into the processes underlying these interactions. This approach can inform practical applications, such as optimizing feeding regimes, and improving animal welfare, ultimately leading to more efficient and sustainable aquaculture systems.

## 7. Declarations

### Declaration of generative AI in scientific writing

Generative AI and AI-assisted technologies were used to improve the readability and language of the manuscript.

#### Acknowledgements

The authors would like to thank the Fishecu team and the Faculty of Fisheries staff for their invaluable assistance during the fieldwork.

## Ethics statement

This study was approved by the Animal Welfare Committee of the Biology Centre CAS according to § 16a of the Act No. 246/1992 Coll., on the protection of animals against cruelty, and Experimental Animal Welfare Commission under the Ministry of Agriculture of the Czech Republic and with permission (Ref. no. MZP/2022/630/2283 1)

## Data archiving statement

The data will be made available on request.

## Author’s contributions

Milan Říha, Lukáš Veselý conceived the ideas and designed methodology; Milan Říha, Lukáš Veselý, Rubén Rabaneda-Bueno, Jaroslav Vrba, Martin Bláha, Vladislav Draštík, Luboš Kočvara collected the data; Milan Říha, Ruben Rabaneda-Bueno, Marie Prchalová, Lenka Kajgrová, Jaroslav Vrba, Trevis B. Meador, Martin Bláha processed data, analyzed samples and analyzed data; Milan Říha led the writing of the manuscript. All authors contributed critically to the drafts and gave final approval for publication.

## References

Adámek, Z., Kajgrová, L., Regenda, J., Roy, K., 2023. Carp pond aquaculture – Concordance of production and nature. Aquaculture Europe 48, 6–15.

Anton-Pardo, M., Adámek, Z., 2015. The role of zooplankton as food in carp pond farming: A review. J. Appl. Ichthyol. 31, 7–14. 10.1111/jai.12852

Anton-Pardo, M., Hlaváč, D., Másílko, J., Hartman, P., Adámek, Z., 2014. Natural diet of mirror and scaly carp (Cyprinus carpio) phenotypes in earth ponds. Folia Zool. 63, 229–237. 10.25225/FOZO.V63.I4.A1.2014

Araujo, G.S., Silva, J.W.A. da, Cotas, J., Pereira, L., 2022. Fish farming techniques: current situation and trends. J. Mar. Sci. Eng. 10, 1598. 10.3390/JMSE10111598

Attia, J., Millot, S., Di-Poï, C., Bégout, M.L., Noble, C., Sanchez-Vazquez, F.J., Terova, G., Saroglia, M., Damsgård, B., 2012. Demand feeding and welfare in farmed fish. Fish Physiol. Biochem. 38, 107–118. 10.1007/S10695-011-9538-4

Bajer, P.G., Lim, H., Travaline, M.J., Miller, B.D., Sorensen, P.W., 2010. Cognitive aspects of food searching behavior in free-ranging wild Common Carp. Environ. Biol. Fishes 88, 295–300. 10.1007/S10641-010-9643-8/METRICS

Balon, E.K., 1995. Origin and domestication of the wild carp, *Cyprinus carpio*: from Roman gourmets to the swimming flowers. Aquaculture 129, 3–48. 10.1016/0044-8486(94)00227-F

Banet, N. V., Fieberg, J., Sorensen, P.W., 2022. Migration, homing and spatial ecology of common carp in interconnected lakes. Ecol. Freshw. Fish 31, 164–176. 10.1111/EFF.12622

Bauer, C., Schlott, G., 2004. Overwintering of farmed common carp (*Cyprinus carpio* L.) in the ponds of a central European aquaculture facility—measurement of activity by radio telemetry. Aquaculture 241, 301–317. 10.1016/J.AQUACULTURE.2004.08.010

Benito, J., Benejam, L., Zamora, L., García-Berthou, E., 2015. Diel cycle and effects of water flow on activity and use of depth by Common carp. Trans. Am. Fish. Soc. 144, 491–501. 10.1080/00028487.2015.1017656

Binhe Gu, Schell, D.M., Xianghao Huang, Fuliang Yie, 1996. Stable isotope evidence for dietary overlap between two planktivorous fishes in aquaculture ponds. Can. J. Fish. Aquat. Sci. 53, 2814–2818. 10.1139/F96-242/ASSET/F96-242.FP.PNG_V03

Brosset, P., Cooke, S.J., Schull, Q., Trenkel, V.M., Soudant, P., Lebigre, C., 2021. Physiological biomarkers and fisheries management. Rev. Fish Biol. Fish. 31, 797–819. 10.1007/S11160-021-09677-5

Bürkner, P.C., 2017. brms: An R package for Bayesian multilevel models using Stan. J. Stat. Softw. 80, 1–28. 10.18637/JSS.V080.I01

Calabrese, J.M., Fleming, C.H., Gurarie, E., 2016. ctmm: an r package for analyzing animal relocation data as a continuous-time stochastic process. Methods Ecol. Evol. 7, 1124–1132. 10.1111/2041-210X.12559

Christensen, R.H.B., 2019. Cumulative link models for ordinal regression with the R package ordinal. Available at: https://cran.r-project.org/web/packages/ordinal/vignettes/clm_article.pdf (accessed 5 June 2020).

Cooke, S.J., Schreer, J.F., 2003. Environmental monitoring using physiological telemetry -A case study examining common carp responses to thermal pollution in a coal-fired generating station effluent. Water. Air. Soil Pollut. 142, 113–136. 10.1023/A:1022082003932/METRICS

Fingerle, A., Larranaga, N., Steingrímsson, S.Ó., 2016. Density-dependent diel activity in stream-dwelling Arctic charr Salvelinus alpinus. Ecol. Evol. 6, 3965–3976. 10.1002/ECE3.2177

Füllner, G., 2015. Traditional feeding of common carp and strategies of replacement of fish meal. In: Biology and Ecology of Carp. Taylor & Francis, Boca Raton, pp. 135–163.

Gamboa-Delgado, J., Molina-Poveda, C., Godínez-Siordia, D.E., Villarreal-Cavazos, D., Ricque-Marie, D., Cruz-Suárez, L.E., 2014. Application of stable isotope analysis to differentiate shrimp extracted by industrial fishing or produced through aquaculture practices. Can. J. Fish. Aquat. Sci. 71, 1520–1528. 10.1139/CJFAS-2014-0005

García-Berthou, E., 2001. Size-and depth-dependent variation in habitat and diet of the common carp (*Cyprinus carpio*). Aquat. Sci. 63, 466–476. 10.1007/S00027-001-8045-6/METRICS

Gelman, A., Carlin, J.B., Stern, H.S., Dunson, D.B., Vehtari, A., Rubin, D.B., 2013. Bayesian data analysis. Chapman and Hall/CRC, New York.

Ghosal, R., Eichmiller, J.J., Witthuhn, B.A., Peter, |, Sorensen, W., Sorensen, P.W., Eichmiller, J., Technical, A., 2018. Attracting Common Carp to a bait site with food reveals strong positive relationships between fish density, feeding activity, environmental DNA, and sex pheromone release that could be used in invasive fish management. Ecol. Evol. 8, 6714–6727. 10.1002/ECE3.4169

Górecki, M.T., Mazurkiewicz, J., Myczko, Ł., 2019. Influence of the presence of ide *Leuciscus idus* on the boldness of common carp *Cyprinus carpio*. Aquaculture 505, 253–255. 10.1016/J.AQUACULTURE.2019.02.069

Hansen, E.A., Closs, G.P., 2005. Diel activity and home range size in relation to food supply in a drift-feeding stream fish. Behav. Ecol. 16, 640–648. 10.1093/beheco/ari036

Hartman, P., Regenda, J., 2016. Praktika v rybníkářství [Practice in fish farming, in Czech]. Jihočeská univerzita v Českých Budějovicích, Fakulta rybářství a ochrany vod.

Hlaváč, D., Adámek, Z., Hartman, P., Másílko, J., 2014. Effects of supplementary feeding in carp ponds on discharge water quality: A review. Aquac. Int. 22, 299–320. 10.1007/S10499-013-9718-6/METRICS

Hundt, P.J., White, L.A., Craft, M.E., Bajer, P.G., 2022. Social associations in common carp (*Cyprinus carpio*): Insights from induced feeding aggregations for targeted management strategies. Ecol. Evol. 12, e8666. 10.1002/ECE3.8666

Huntingford, F.A., Andrew, G., Mackenzie, S., Morera, D., Coyle, S.M., Pilarczyk, M., Kadri, S., 2010. Coping strategies in a strongly schooling fish, the common carp *Cyprinus carpio*. J. Fish Biol. 76, 1576–1591. 10.1111/J.1095-8649.2010.02582.X

Jurajda, P., Adámek, Z., Roche, K., Mrkvová, M., Štarhová, D., Prášek, V., Zukal, J., 2016. Carp feeding activity and habitat utilisation in relation to supplementary feeding in a semi-intensive aquaculture pond. Aquac. Int. 24, 1627–1640. 10.1007/s10499-016-0061-6

Kajgrova, L., Adamek, Z., Regenda, J., Bauer, C., Stejskal, V., Pecha, O., Hlavac, D., 2021. Macrozoobenthos assemblage patterns in European carp (*Cyprinus carpio*) ponds − the importance of emersed macrophyte beds. Knowl. Manag. Aquat. Ecosyst. 2020-January, 9. 10.1051/KMAE/2021008

Kajgrová, L., Blabolil, P., Drozd, B., Roy, K., Regenda, J., Šorf, M., Vrba, J., 2022. Negative effects of undesirable fish on common carp production and overall structure and functioning of fishpond ecosystems. Aquaculture 549, 737811. 10.1016/J.AQUACULTURE.2021.737811

Kajgrová, L., Kolar, V., Roy, K., Adámek, Z., Blabolil, P., Kopp, R., Mráz, J., Musil, M., Pecha, O., Pechar, L., Potužák, J., Vrba, J., 2024. A stoichiometric insight into the seasonal imbalance of phosphorus and nitrogen in central European fishponds. Environ. Sci. Eur. 36, 1–9. 10.1186/S12302-024-00968-9/FIGURES/3

Klaren, P.H.M., van Dalen, S.C.M., Atsma, W., Spanings, F.A.T., Hendriks, J., Flik, G., 2013. Voluntary timing of food intake increases weight gain and reduces basal plasma cortisol levels in common carp (*Cyprinus carpio* L.). Physiol. Behav. 122, 120–128. 10.1016/J.PHYSBEH.2013.08.020

Kolářová, J., Velíšek, J., Nepejchalová, L., Svobodová, J., Kouřil, J., Hamáčková, J., Máchová, J., Plačková, V., Hajšlová, J., Holadová, K., Kocourek, V., Elimánková, E., Modrá, H., Dobšíková, R., Groch, L., Novotný, L., 2007. Anestetika pro ryby [Anestitics for fish, in Czech]. Jihočeská univerzita v Českých Budějovicích, Fakulta rybářství a ochrany vod, Vodňany, 6–13. ISBN 978-80-85887-61-7.

Lammens, E.H.R.R., Hoogenboezem, W., 1991. Diets and feeding behaviour. In: Winfield, I.J., Nelson, J.S. (Eds.), Cyprinid Fishes: Systematics, Biology and Exploitation. Springer Netherlands, Dordrecht, pp. 353–376.

Lennox, R.J., Adam, T., Riha, M., Klappstein, N., Monk, C.T., Vollset, K.W., Beumer, L.T., 2024. Movement in 3D: Novel opportunities for understanding animal behaviour and space use. Ethology 0, e13529. 10.1111/ETH.13529

Martin-Creuzburg, D., Von Elert, E., Hoffmann, K.H., 2008. Nutritional constraints at the cyanobacteria— Daphnia magna interface: The role of sterols. Limnol. Oceanogr. 53, 456–468. 10.4319/LO.2008.53.2.0456

Martínez Del Rio, C., Wolf, N., Carleton, S.A., Gannes, L.Z., 2009. Isotopic ecology ten years after a call for more laboratory experiments. Biol. Rev. 84, 91–111. 10.1111/J.1469-185X.2008.00064.X

Matte, J.M.O., Fraser, D.J., Grant, J.W.A., 2021. Mechanisms of density dependence in juvenile salmonids: prey depletion, interference competition, or energy expenditure? Ecosphere 12, e03567. 10.1002/ECS2.3567

Mccutchan, James H, Lewis, William M, Kendall, C., Mcgrath Mccutchan, C.C., Lewis, J.H., Kendall, W.M., Mcgrath, C., Mccutchan, J H, Lewis, W M, Mcgrath, C.C., 2003. Variation in trophic shift for stable isotope ratios of carbon, nitrogen, and sulfur. Oikos 102, 378–390. 10.1034/J.1600-0706.2003.12098.X

Mehner, T., Rapp, T., Monk, C.T., Beck, M.E., Trudeau, A., Kiljunen, M., Hilt, S., Arlinghaus, R., 2019. Feeding aquatic ecosystems: Whole-lake experimental addition of angler’s ground bait strongly affects omnivorous fish despite low contribution to lake carbon budget. Ecosystems 22, 346–362. 10.1007/S10021-018-0273-X/METRICS

Monk, C.T., Arlinghaus, R., 2017. Encountering a bait is necessary but insufficient to explain individual variability in vulnerability to angling in two freshwater benthivorous fish in the wild. PLoS One 12, e0173989. 10.1371/JOURNAL.PONE.0173989

Monk, C.T., Chéret, B., Czapla, P., Hühn, D., Klefoth, T., Eschbach, E., Hagemann, R., Arlinghaus, R., 2020. Behavioural and fitness effects of translocation to a novel environment: wholeLJlake experiments in two aquatic top predators. J. Anim. Ecol. 1365–2656.13298. 10.1111/1365-2656.13298

Moore, J.W., Semmens, B.X., 2008. Incorporating uncertainty and prior information into stable isotope mixing models. Ecol. Lett. 11, 470–480. 10.1111/J.1461-0248.2008.01163.X

Musil, M., Novotná, K., Potužák, J., Hůda, J., Pechar, L., 2015. Impact of topmouth gudgeon (Pseudorasbora parva) on production of common carp (*Cyprinus carpio*) — question of natural food structure. Biologia (Bratisl). 69, 1757–1769. 10.2478/S11756-014-0483-4

Nathan, R., Monk, C.T., Arlinghaus, R., Adam, T., Alós, J., Assaf, M., Baktoft, H., Beardsworth, C.E., Bertram, M.G., Bijleveld, A.I., Brodin, T., Brooks, J.L., Campos-Candela, A., Cooke, S.J., Gjelland, K., Gupte, P.R., Harel, R., Hellström, G., Jeltsch, F., Killen, S.S., Klefoth, T., Langrock, R., Lennox, R.J., Lourie, E., Madden, J.R., Orchan, Y., Pauwels, I.S., Říha, M., Roeleke, M., Schlägel, U.E., Shohami, D., Signer, J., Toledo, S., Vilk, O., Westrelin, S., Whiteside, M.A., Jarić, I., 2022. Big-data approaches lead to an increased understanding of the ecology of animal movement. Science (80-.). 375. 10.1126/SCIENCE.ABG1780/SUPPL_FILE/SCIENCE.ABG1780_MOVIES_S1_TO_S5. ZIP

Nordahl, O., Tibblin, P., Koch-Schmidt, P., Berggren, H., Larsson, P., Forsman, A., 2018. Sun-basking fish benefit from body temperatures that are higher than ambient water. Proc. R. Soc. B Biol. Sci. 285. 10.1098/RSPB.2018.0639

NRC (National Research Council), 2011. Nutrient requirements of fish and shrimp. National Academy Press, Washington, DC, 70 p.

Orpwood, J.E., Griffiths, S.W., Armstrong, J.D., 2006. Effects of food availability on temporal activity patterns and growth of Atlantic salmon. J. Anim. Ecol. 75, 677–685. 10.1111/J.1365-2656.2006.01088.X

Orság, M., Meitner, J., Fischer, M., Svobodová, E., Kopp, R., Mareš, J., Spurný, P., Pechar, L., Beděrková, I., Hanuš, J., Semerádová, D., Balek, J., Radojičić, M., Hanel, M., Vizina, A., Žalud, Z., Trnka, M., 2023. Estimating Heat Stress Effects on the Sustainability of Traditional Freshwater Pond Fishery Systems under Climate Change. Water 2023, Vol. 15, Page 1523 15, 1523. 10.3390/W15081523

Özgül, A., Birnie-Gauvin, K., Abecasis, D., Alós, J., Aarestrup, K., Reubens, J., Bolland, J., Lök, A., Edwards, J.E., Pengal, P., Prchalová, M., Říha, M., Pickholtz, R., Vollset, K.W., Afonso, P., Davidsen, J.G., Arlinghaus, R., Ünal, V., Lennox, R.J., 2024. Tracking aquatic animals for fisheries management in European waters. Fish. Manag. Ecol. 31, e12706. 10.1111/FME.12706

Parata, L., Sammut, J., Egan, S., 2021. Opportunities for microbiome research to enhance farmed freshwater fish quality and production. Rev. Aquac. 13, 2027–2037. 10.1111/RAQ.12556

Pechar, L., 2000. Impacts of long-term changes in fishery management on the trophic level water quality in Czech fish ponds. Fish. Manag. Ecol. 7, 23–31. 10.1046/J.1365-2400.2000.00193.X

Post, D.M., 2002. Using stable isotopes to estimate trophic position: Models, methods, and assumptions. Ecology 83, 703–718. 10.1890/0012-9658(2002)083[0703:USITET]2.0.CO;2

Potužák, J., Hůda, J., Pechar, L., 2007. Changes in fish production effectivity in eutrophic fishponds - Impact of zooplankton structure. Aquac. Int. 15, 201–210. 10.1007/S10499-007-9085-2/METRICS

R Development Core Team, R., 2024. R: a language and environment for statistical computing. R Found. Stat. Comput., R Foundation for Statistical Computing. 10.1007/978-3-540-74686-7

Rahman, M.M., 2015. Role of common carp (*Cyprinus carpio*) in aquaculture production systems. Front. Life Sci. 8, 399–410. 10.1080/21553769.2015.1045629

Rahman, M.M., Meyer, C.G., 2009. Effects of food type on diel behaviours of common carp *Cyprinus carpio* in simulated aquaculture pond conditions. J. Fish Biol. 74, 2269–78. 10.1111/j.1095-8649.2009.02236.x

Ray, A.J., Drury, T.H., Cecil, A., 2017. Comparing clear-water RAS and biofloc systems: Shrimp (*Litopenaeus vannamei)* production, water quality, and biofloc nutritional contributions estimated using stable isotopes. Aquac. Eng. 77, 9–14. 10.1016/J.AQUAENG.2017.02.002

Rigby, R.A., Stasinopoulos, D.M., 2009. A Flexible Regression Approach Using GAMLSS in R. London Metropolitan University, London, UK.

Říha, M., Gjelland, K., Děd, V., Eloranta, A.P., Rabaneda-Bueno, R., Baktoft, H., Vejřík, L., Vejříková, I., Draštík, V., Šmejkal, M., Holubová, M., Jůza, T., Rosten, C., Sajdlová, Z., Økland, F., Peterka, J., 2021. Contrasting structural complexity differentiate hunting strategy in an ambush apex predator. Sci. Rep. 11, 17472. 10.1038/s41598-021-96908-1

Říha, M., Rabaneda-Bueno, R., Jarić, I., Souza, A.T., Vejřík, L., Draštík, V., Blabolil, P., Holubová, M., Jůza, T., Gjelland, K., Rychtecký, P., Sajdlová, Z., Kočvara, L., Tušer, M., Prchalová, M., Seďa, J., Peterka, J., 2022. Seasonal habitat use of three predatory fishes in a freshwater ecosystem. Hydrobiologia 849, 3351–3371. 10.1007/S10750-022-04938-1

Říha, M., Rabaneda-Bueno, R., Vejřík, L., Jarić, I., Prchalová, M., Šmejkal, M., Čech, M., Draštík, V., Blabolil, P., Holubová, M., Jůza, T., Gjelland, K.Ø., Sajdlová, Z., Kočvara, L., Tušer, M., Peterka, J., 2024. Hungry catfish – effect of prey availability on movement dynamics of a top predator. Freshw. Biol.

Roy, K., Kajgrova, L., Capkova, L., Zabransky, L., Petraskova, E., Dvorak, P., Nahlik, V., Kuebutornye, F.K.A., Blabolil, P., Blaha, M., Vrba, J., Mraz, J., 2024. Synergistic digestibility effect by planktonic natural food and habitat renders high digestion efficiency in agastric aquatic consumers. Sci. Total Environ. 927, 172105. 10.1016/J.SCITOTENV.2024.172105

Roy, K., Másílko, J., Kajgrova, L., Kuebutornye, F.K.A., Oberle, M., Mraz, J., 2023. End-of-season supplementary feeding in European carp ponds with appropriate plant protein and carbohydrate combinations to ecologically boost productivity: Lupine, rapeseed and, triticale. Aquaculture 577, 739906. 10.1016/J.AQUACULTURE.2023.739906

Roy, K., Vrba, J., Kajgrova, L., Mraz, J., 2022. The concept of balanced fish nutrition in temperate European fishponds to tackle eutrophication. J. Clean. Prod. 364, 132584. 10.1016/J.JCLEPRO.2022.132584

Roy, K., Vrba, J., Kaushik, S.J., Mraz, J., 2020. Nutrient footprint and ecosystem services of carp production in European fishponds in contrast to EU crop and livestock sectors. J. Clean. Prod. 270, 122268. 10.1016/J.JCLEPRO.2020.122268

Schälicke, S., Sobisch, L.Y., Martin-Creuzburg, D., Wacker, A., 2019. Food quantity–quality co-limitation: Interactive effects of dietary carbon and essential lipid supply on population growth of a freshwater rotifer. Freshw. Biol. 64, 903–912. 10.1111/FWB.13272

Sibbing, F.A., Osse, J.W.M., Terlouw, A., 1986. Food handling in the carp (*Cyprinus carpio*): its movement patterns, mechanisms and limitations. J. Zool. 210, 161–203. 10.1111/J.1469-7998.1986.TB03629.X

Sogard, S.M., Olla, B.L., 1996. Food deprivation affects vertical distribution and activity of a marine fish in a thermal gradient: potential energy-conserving mechanisms. Mar. Ecol. Prog. Ser. 133, 43–55. 10.3354/MEPS133043

Stanivuk, J., Berzi-Nagy, L., Gyalog, G., Ardó, L., Vitál, Z., Plavša, N., Krstović, S., Fazekas, G.L., Horváth, Á., Ljubobratović, U., 2024. The rank of intensification factors strength in intensive pond production of common carp (*Cyprinus carpio* L.). Aquaculture 583, 740584. 10.1016/J.AQUACULTURE.2024.740584

Stasinopoulos, D.M., Rigby, R.A., 2007. Generalized additive models for Location Scale and Shape (GAMLSS) in R. 10.18637/jss.v023.i07

Stock, B.C., Semmens, B.X., 2016. Unifying error structures in commonly used biotracer mixing models. Ecology 97, 2562–2569. 10.1002/ECY.1517

Vejřík, L., Matějíčková, I., Jůza, T., Frouzová, J., Seďa, J., Blabolil, P., Ricard, D., Vašek, M., Kubečka, J., Říha, M., Čech, M., 2016. Small fish use the hypoxic pelagic zone as a refuge from predators. Freshw. Biol. 61. 10.1111/fwb.12753

Verdegem, M., Buschmann, A.H., Latt, U.W., Dalsgaard, A.J.T., Lovatelli, A., 2023. The contribution of aquaculture systems to global aquaculture production. J. World Aquac. Soc. 54, 206–250. 10.1111/JWAS.12963

Vrba, J., Šorf, M., Nedoma, J., Benedová, Z., Kröpfelová, L., Šulcová, J., Tesařová, B., Musil, M., Pechar, L., Potužák, J., Regenda, J., Šimek, K., Řeháková, K., 2024. Top-down and bottom-up control of plankton structure and dynamics in hypertrophic fishponds. Hydrobiologia 851, 1095–1111. 10.1007/S10750-023-05312-5/TABLES/4

Watkinson, D.A., Charles, C., Enders, E.C., 2021. Spatial ecology of common carp (*Cyprinus carpio*) in Lake Winnipeg and its potential for management actions. J. Great Lakes Res. 47, 583–591. 10.1016/J.JGLR.2021.03.004

Weber, M.J., Brown, M.L., 2009. Effects of Common Carp on Aquatic Ecosystems 80 Years after “Carp as a Dominant”: Ecological Insights for Fisheries Management. Rev. Fish. Sci. 17. 10.1080/10641260903189243

Žák, J., 2021. Diel pattern in common carp landings from angling competitions corresponds to their assumed foraging activity. Fish. Res. 243, 106086. 10.1016/J.FISHRES.2021.106086

Zanden, M.J. Vander, Rasmussen, J.B., 2001. Variation in δ15N and δ13C trophic fractionation: Implications for aquatic food web studies. Limnol. Oceanogr. 46, 2061–2066. 10.4319/LO.2001.46.8.2061

Zhou, B.S., Wu, R.S.S., Randall, D.J., Lam, P.K.S., Ip, Y.K., Chew, S.F., 2000. Metabolic adjustments in the common carp during prolonged hypoxia. J. Fish Biol. 57, 1160–1171. 10.1111/J.1095-8649.2000.TB00478.X

Zion, B., Barki, A., Grinshpon, J., Rosenfeld, L., Karplus, I., 2007. Social facilitation of acoustic training in the common carp *Cyprinus carpio* (L.). Behaviour 144, 611–630. 10.1163/156853907781347781

