## Supplementary materials 1 for "A fresh perspective on carp feeding behavior in an aquaculture pond and its consequences for individual growth"

#### Testing of the telemetry system performance and accuracy

### The system's localization ability and accuracy were evaluated using a series of stationary reference tags (MM-M-16-50-TP, burst rate: 25 seconds, Lotek Wireless Inc., Canada) installed at predetermined positions (5–6 tags) across open water, near-bottom, and littoral zones, along with a series of mobile tags (4 tags) towed behind a boat equipped with a GPS device (Garmin GPSMAP 60C). Stationary tags remained deployed throughout the entire system installation period, while mobile testing was conducted in June 2022 and October 2023. These efforts generated two datasets for assessing the positioning system's accuracy: stationary and mobile tests.

#### Stationary reference tags

In 2022, a total of 2.31 million locations were detected from stationary tags out of 3.82 million transmitted pings, indicating that 61% of transmitted pings were successfully detected and used for location calculations. Detection rates were lower near the bottom (30%–50%) compared to the littoral and open water zones (60%–80%). Despite this variation, the distance error and error variability were consistent across all habitats, demonstrating very high precision and low variability. Nearly all locations were detected with errors ranging from 1 to 2.5 meters from the true location, with the standard deviation (SD) of error ranging from 2 to 7 meters (Fig. 1 SM).

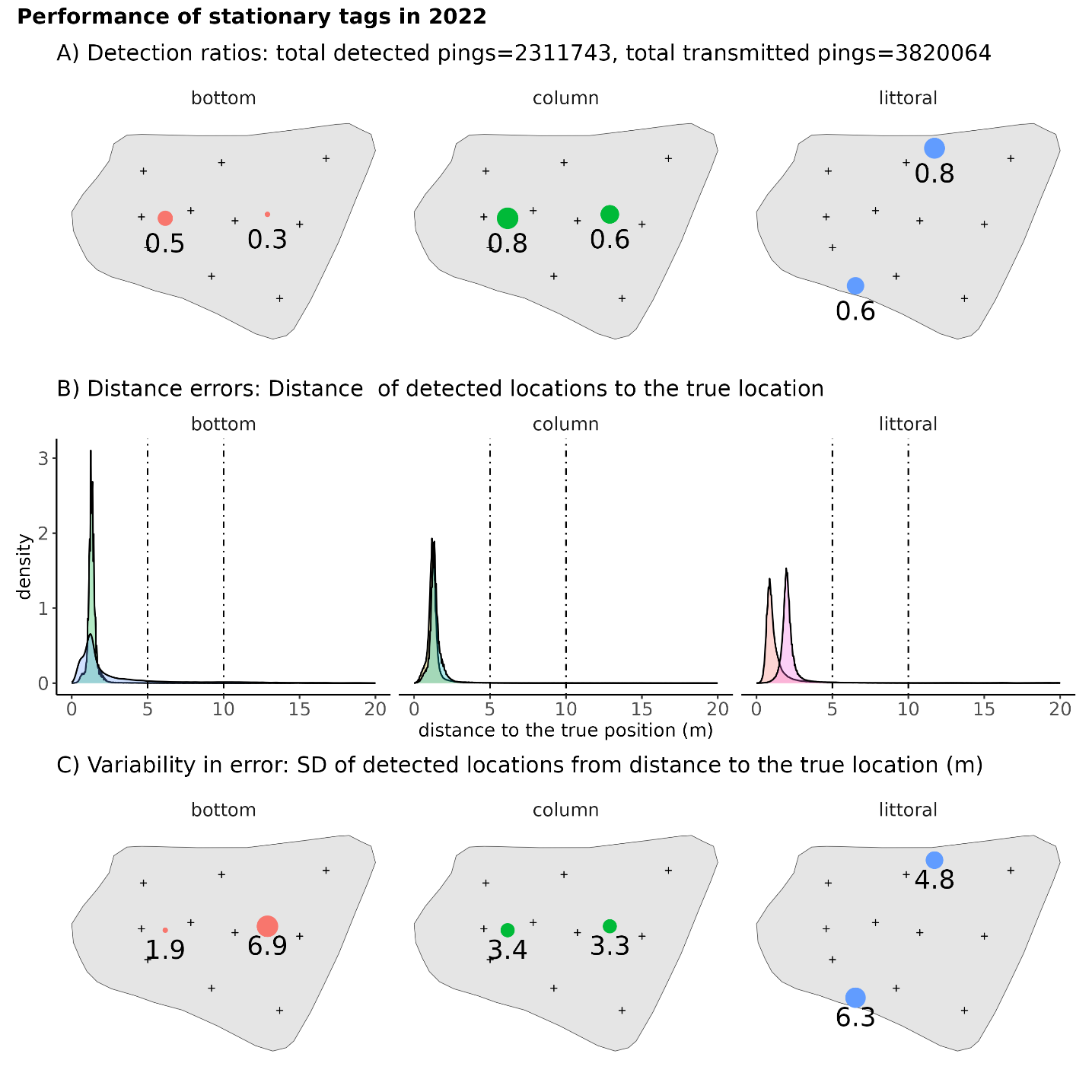

**Figure 1 SM1**: Performance of stationary tags in 2022. **A)** Detection ratios: Proportion of detected pings relative to total transmitted pings for stationary tags in different habitats. **B)** Distance errors: Distribution of errors in detected tag locations relative to their true positions. **C)** Variability in error: Standard deviation (SD) of distance errors for detected locations.

In 2023, a total of 2.29 million locations were detected from stationary tags out of 3.23 million transmitted pings, indicating that 71% of transmitted pings were successfully detected and used for location calculations. A similar trend was observed as in 2022, with detection rates being lower near the bottom (40%) compared to the littoral and open water zones (70%–90%). Precision and variability remained consistent across most tags, except for one littoral tag, which exhibited a higher location error (9.3 m) and another with higher temporal variability (SD = 9.6 m; Fig. 2 SM).

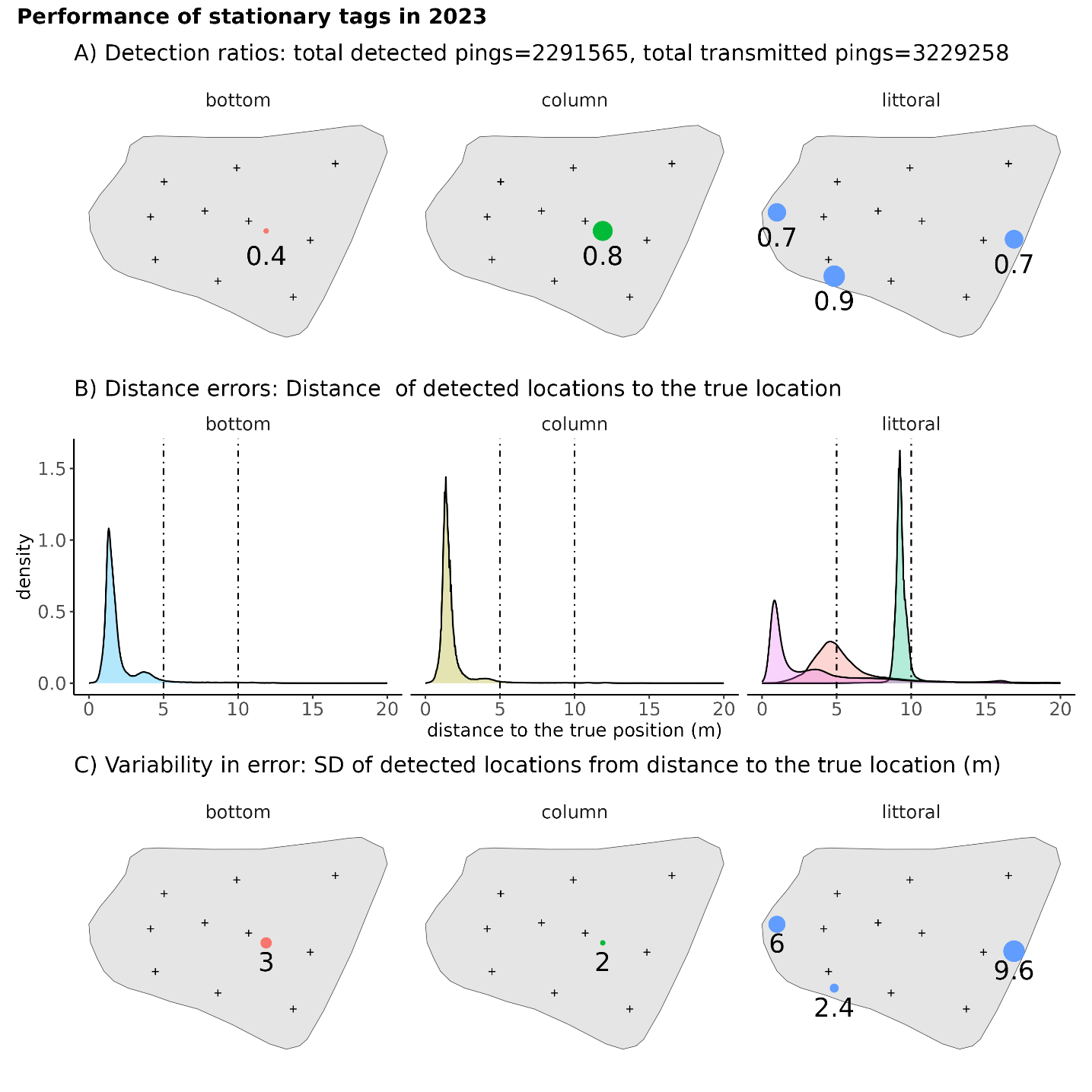

**Figure 2 SM**: Performance of stationary tags in 2023. **A)** Detection ratios: Proportion of detected pings relative to total transmitted pings for stationary tags in different habitats. **B)** Distance errors: Distribution of errors in detected tag locations relative to their true positions. **C)** Variability in error: Standard deviation (SD) of distance errors for detected locations.

#### Mobile test

In 2022, a total of 917 locations were detected from stationary tags out of 1,132 transmitted pings, resulting in an 81% detection rate, indicating that the majority of transmitted pings were successfully detected and used for location calculations. The detection ratio, precision, and variability in precision were consistent across all dragged tags. Detection ratios ranged from 80% to 82%, precision ranged from 6.6 to 6.8 meters, and standard deviation (SD) ranged from 3.2 to 6.3 meters. Precision was uniform throughout the pond, with no areas showing significantly poorer coverage (Fig. 3 SM).

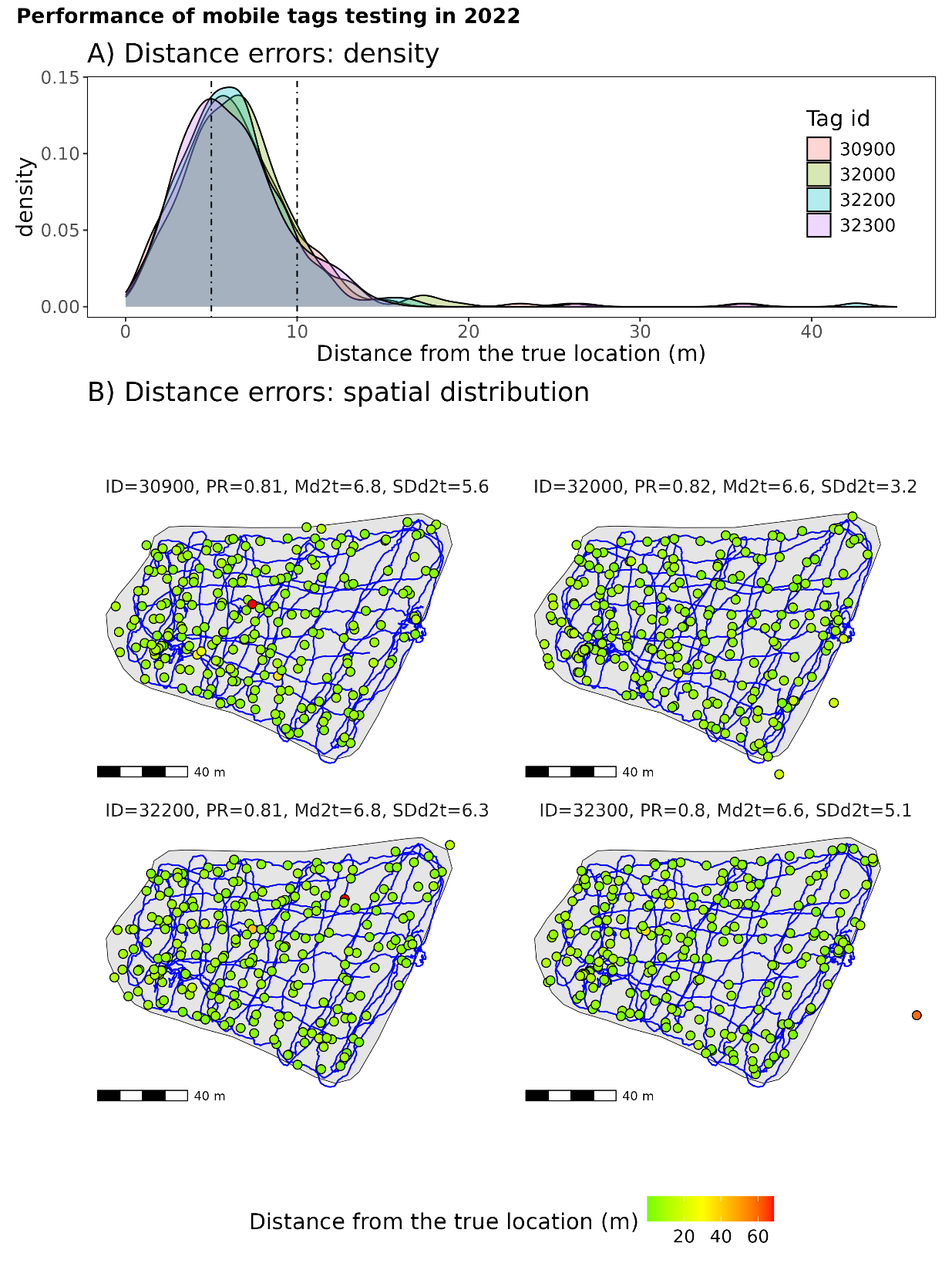

**Figure 3: Performance of mobile tags testing in 2023. A)** Density distribution of errors in the detected locations relative to the true locations of the mobile tags. The median distance errors (Md2t) ranged from 4.0 to 4.8 meters, indicating improved precision compared to 2022. **B)** Spatial distribution of detected locations for each mobile tag (Tag IDs: 30900, 32300, 32500, 32900) overlaid on the boat's trajectory (blue line). The proportion of detected signals (PR) ranged from 0.80 to 0.88, with variability in precision (SDd2t) ranging from 3.3 to 6.7 meters. The color scale represents the distance of each detected location from the true location, highlighting spatial variations in accuracy. The blue line indicates the trajectory of the boat towing the reference tags.

In 2023, a total of 573 locations were detected from stationary tags out of 682 transmitted pings, resulting in an 84% detection rate, indicating that the majority of transmitted pings were successfully detected and converted into location data. The detection ratio, precision, and variability in precision remained consistent across all dragged tags. Detection ratios ranged from 80% to 88%, precision ranged from 4.0 to 4.8 meters, and the standard deviation (SD) ranged from 3.3 to 6.7 meters. Precision was uniformly distributed throughout the pond, with no areas exhibiting significantly poorer coverage (Fig. 4 SM).

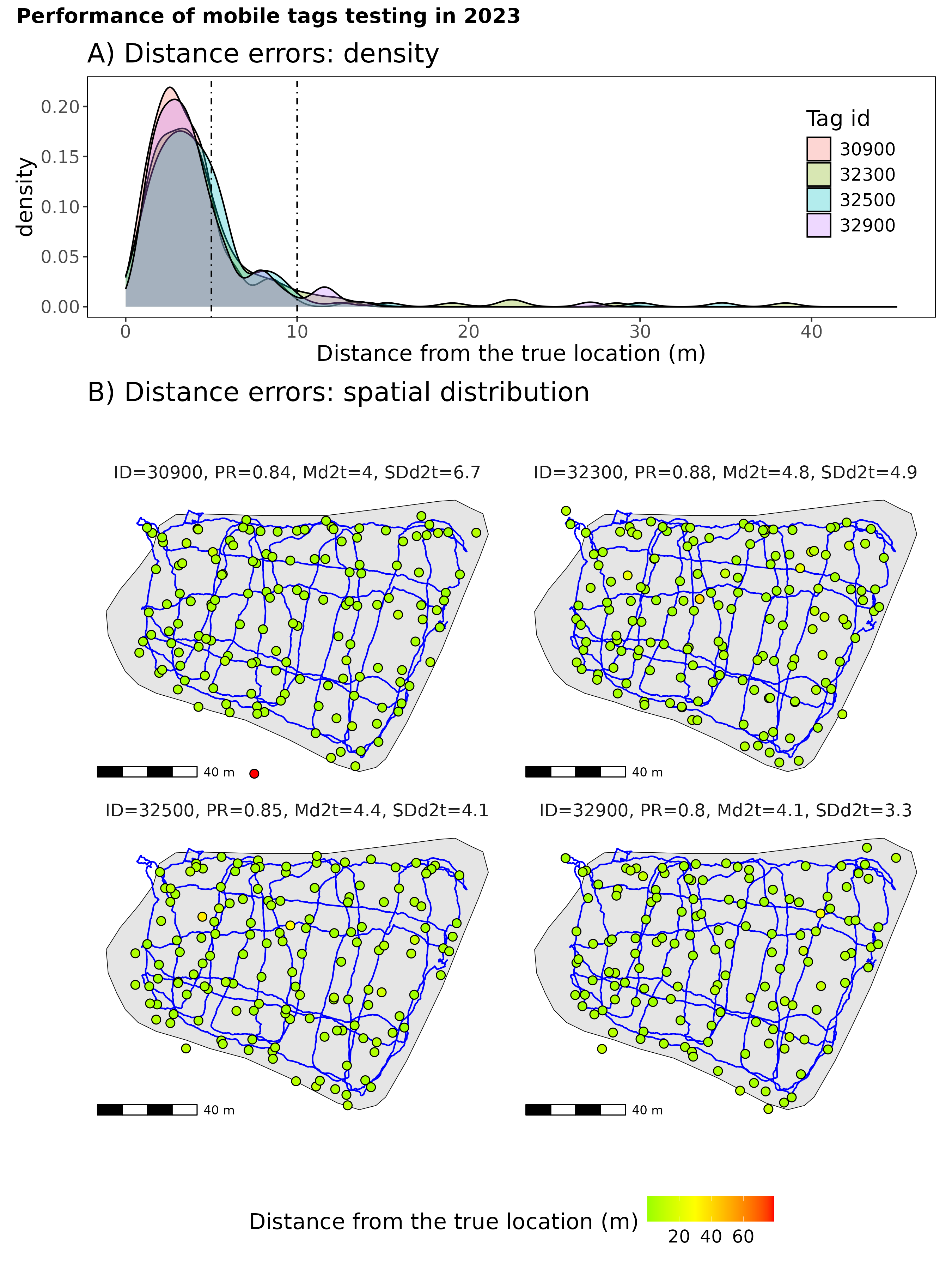

**Figure 4: Performance of mobile tags testing in 2023. A)** Density distribution of errors in the detected locations relative to the true locations of the mobile tags. The median distance errors (Md2t) ranged from 4.0 to 4.8 meters. **B)** Distance errors: Spatial distribution of detected locations for each mobile tag (Tag IDs: 30900, 32300, 32500, 32900) overlaid on the boat's trajectory (blue line). The proportion of detected signals (PR) ranged from 0.80 to 0.88, with variability in precision (SDd2t) ranging from 3.3 to 6.7 meters. The color scale represents the distance of each detected location from the true location, highlighting spatial variations in accuracy. The blue line indicates the trajectory of the boat towing the reference tags.

#### Summary

Across both years, the system demonstrated reliability, with high detection rates and consistent spatial precision, ensuring robust performance of the telemetry system and obtained data for our study. The system maintained high detection rates. Precision and error variability were very consistent across different areas of the pond, ensuring even spatial coverage without significant gaps. These results confirm the system's capability for reliable localization of telemetry tags, supporting its effectiveness for tracking and monitoring aquatic systems.
