## Supplementary materials 2 for "A fresh perspective on carp feeding behavior in an aquaculture pond and its consequences for individual growth"

Individual space use:

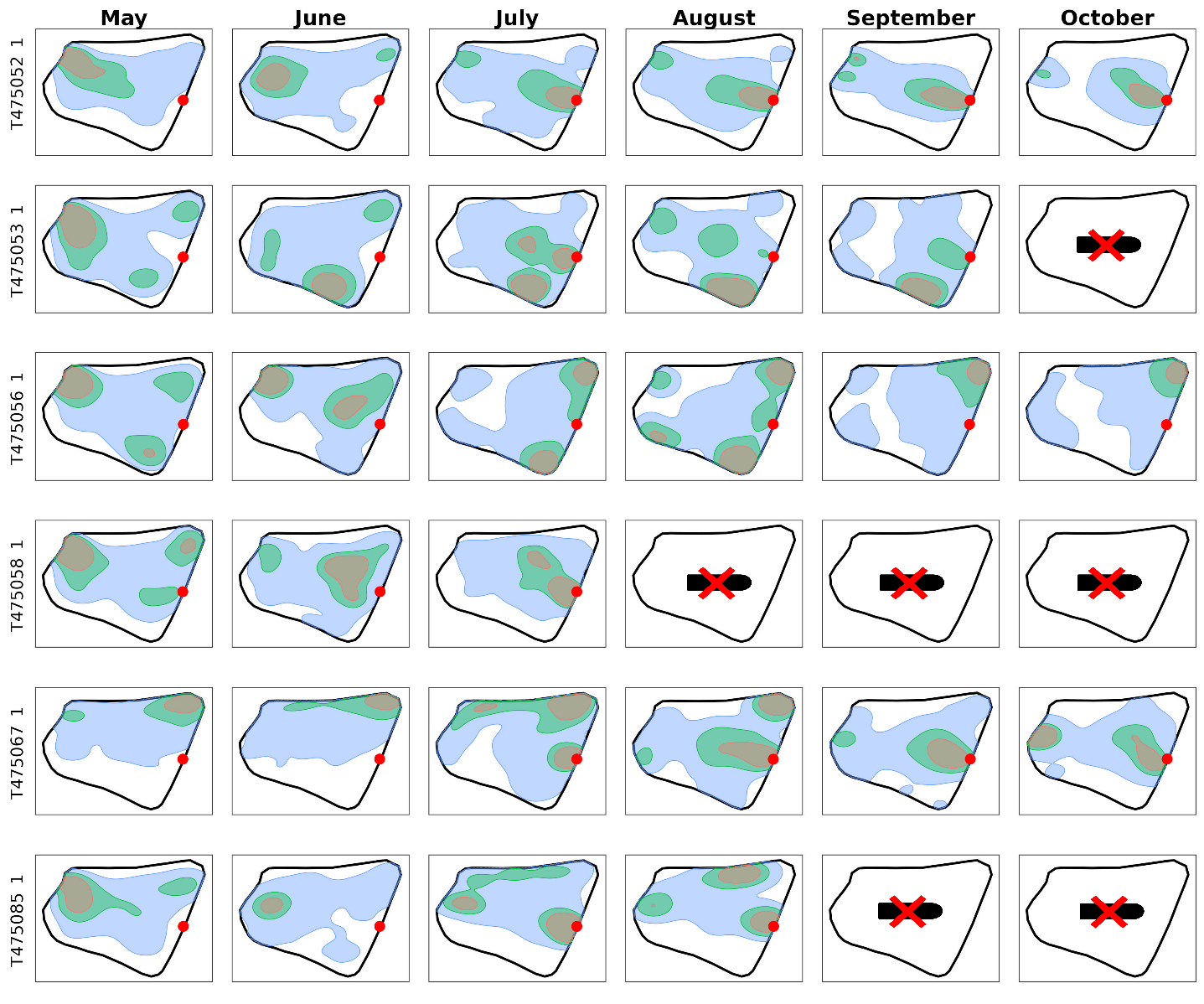

**Figure 1 SM2.** Spatiotemporal visualization of the activity centers of all tracked individuals (population kernel density estimation) across months for 2022. The kernel density estimates represent utilization distributions (UD) of tracked individuals, with gray indicating 25% UD, green 50% UD, and blue 95% UD. The red dot marks the designated feeding ground. Each panel represents a month, lines represents individuals. Pictograms indicate months when tags malfunctioned, leading to missing data.

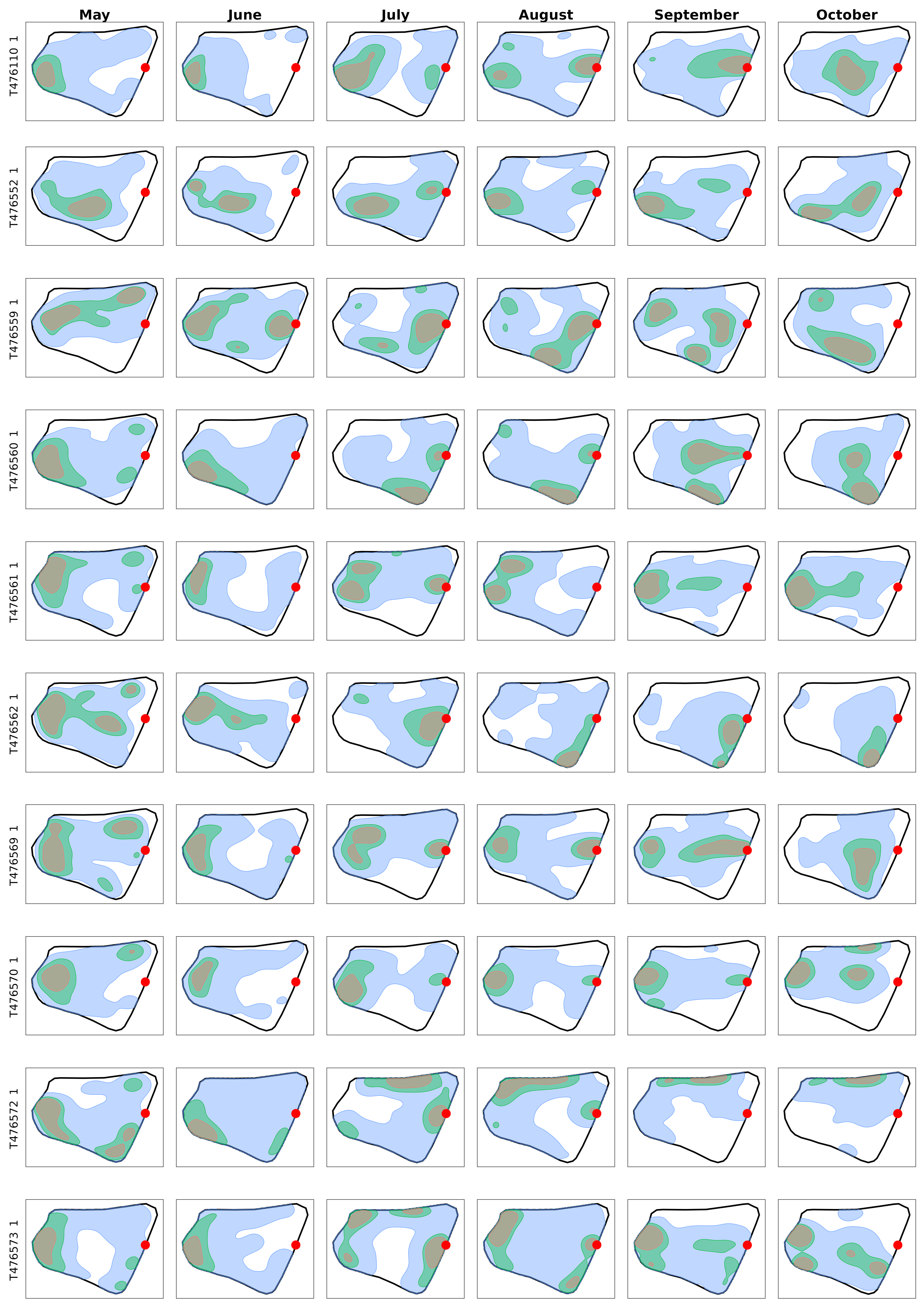

**Figure 2 SM2.** Spatiotemporal visualization of the activity centers of all tracked individuals (population kernel density estimation) across months for 2023. The kernel density estimates represent utilization distributions (UD) of tracked individuals, with gray indicating 25% UD, green 50% UD, and blue 95% UD. The red dot marks the designated feeding ground. Each panel represents a month, lines represents individuals.
